# NAD+ boosting by oral nicotinamide mononucleotide administration regulates key metabolic and immune pathways through SIRT1 dependent and independent mechanisms to mitigate diet-induced obesity and dyslipidemia in mice

**DOI:** 10.1101/2025.06.20.660826

**Authors:** Yasser Majeed, Najeeb M. Halabi, Rudolf Engelke, Hina Sarwath, Muna N. Al-Noubi, Sunkyu Choi, Aisha Al-Malki, Maha V. Agha, Muneera Vakayil, Lotfi Chouchane, Frank Schmidt, Nayef A. Mazloum

## Abstract

Sirtuins are NAD+-dependent histone deacetylases that play a key role in metabolism. Sirtuin activity is compromised in aging and metabolic disorders, and pharmacological strategies that promote sirtuin function including NAD+ boosting approaches show potential as therapeutics. To study the impact of nicotinamide mononucleotide (NMN) supplementation in mice in high fat diet (HFD) induced obesity and the role of SIRT1, a sirtuin family member, in mediating the NMN response, we administered NMN to mice in drinking water to boost NAD+ in control and in inducible SIRT1 knock-out mouse models and performed a combination of metabolic phenotyping, lipid profiling and plasma proteomics in these mice. We discovered that supplementation with NMN mitigated diet induced weight gain by enhancing energy expenditure, corrected dyslipidemia, and reversed perturbations in fasting blood glucose, all in a SIRT1-dependent manner. On the other hand, NMN-induced reductions in fat mass, fluid mass, eWAT and mesenteric WAT were SIRT1 independent. Proteomic approaches in plasma samples using O-Link and mass-spectrometry provided novel insights into key obesity- and NMN-dependent changes in circulating molecules with potential relevance to inflammation, liver function, and dyslipidemia. We discovered SIRT1 dependent and independent alterations in key circulating plasma proteins and identified key metabolic and molecular pathways that were significantly affected by HFD, several of which were reverted by oral NMN administration. Glucose metabolism, cholesterol metabolism and immune-related pathways are among the most significantly affected changes. Causal analysis of proteomic data suggests that observed effects could be mediated by transcription regulators FBXW7, ADIPOR2 and PRDM16. Collectively, our data supports the hypothesis that promoting SIRT1 function by boosting NAD+ levels in vivo may be a useful strategy to mitigate obesity and associated cardiovascular complications such as dyslipidemia.

## Introduction

Statistics from the World Health Organization show that the prevalence of obesity – defined by a body mass index ≥ 30 kg/m^2^ - has significantly increased in the last five decades, and that obese subjects are at a significantly higher risk of morbidity and mortality from obesity-related complications including cardiovascular disease (CVD) [1, 2]. Adipose tissue (AT) regulates an array of key metabolic processes, including storage of lipids, regulation of body temperature, and secretion of hormones like leptin and adiponectin [3, 4]. The most abundant cell-type in AT is the adipocyte, but other AT-resident cell-types with functional importance include fibroblasts, macrophages, and adipocyte progenitors, which respond to metabolic demands by terminally differentiating into adipocytes [5]. In contrast, AT is dysfunctional in obesity and is characterized by a hypoxic environment, inflammation, adipocyte hypertrophy, adipocyte loss due to apoptosis, cellular senescence, and immune cell infiltration, all of which combine to disrupt organ function and contribute to metabolic syndrome – an umbrella term that includes type 2 diabetes (T2D), hypertension, dyslipidemia, and non-alcoholic fatty liver disease (NAFLD) [6–10]. Obese AT also secretes pro-inflammatory cytokines like interleukin-6 (IL-6), tumor necrosis factor-α (TNF-α), and serum amyloid A3 (SAA3) into the circulation [11], and these factors are known to dampen insulin-sensitivity and sustain inflammation [12, 13]. Important insights into the systemic changes driven by obesity were obtained by applying plasma proteomics approaches to mouse samples, which identified changes in proteins such as thyroxine-binding globulin, apolipoprotein C-II, apolipoprotein E and A4, fructose-bisphosphate aldolase B, and components of the complement cascade [14]. Changes in the plasma levels of many of these proteins correlated with obesity-associated dyslipidemia and liver dysfunction [15]. Application of O-link proteomics to human samples identified molecules whose plasma levels strongly correlated with body mass index (BMI), visceral adiposity, and weight-loss, including interleukin-6 (IL-6), leptin, fatty acid binding protein 4 (FABP-4), hepatocyte growth factor (HGF), and tumor necrosis factor receptor superfamily (TNFRSF) members 10C, 11A, and 14 [16, 17].

Sirtuin 1 (SIRT1) – a NAD^+^-dependent enzyme that removes acetyl groups from target proteins to generate O-acyl-ADP-ribose and nicotinamide – regulates key metabolic functions by controlling the activity of targets like PPARγ and PGC-1α [18–23]. Data support a role for SIRT1 in promoting mitochondrial biogenesis in adipocytes [24], regulating preadipocyte proliferative potential [25], improving insulin-sensitivity and inflammation [26], and suppressing lipid accumulation [27]. SIRT1 activity also improves skeletal muscle insulin-sensitivity observed during calorie-restriction [28], mitigates AT inflammation [29], and protects against liver steatosis and inflammation [30]. A positive correlation between adipose SIRT1 expression and energy expenditure or insulin-sensitivity has also been reported in human subjects [31]. SIRT1 function depends on NAD^+^, which can be generated from nicotinamide mononucleotide (NMN) by nicotinamide mononucleotide adenylyltransferases (NMNATs) [32]. Therefore, downregulation of these enzymes in the AT of obese mice limits SIRT1 function to promote metabolic dysfunction [29]. Importantly, several studies have demonstrated a decline in cellular NAD^+^ levels during aging and in metabolic disorders [33–35] and strategies that replenished NAD^+^ in mice by supplementation with precursors like NMN and Nicotinamide Riboside (NR) mitigated age-related dysfunction and protected against obesity [36–39]. On the other hand, recent studies in obese mice revealed that exercise-induced metabolic benefits were *reversed* by NMN administration [40], NR administration only mildly improved metabolic function and energy metabolism [41–43], and boosting NAD^+^ levels *in vivo* exacerbated inflammation and cancer progression [44]. Therefore, further investigations are necessary to clarify the effects of NAD^+^ boosting in an obesity context and to establish whether SIRT1 activation contributes to any potential benefits.

Here we examined the impact of dietary NMN supplementation on weight gain, energy expenditure, dyslipidemia, glucose metabolism and plasma proteome in control and SIRT1 KO mice fed chow or high-fat diet (HFD). We discovered that oral administration of NMN in mice significantly mitigated high-fat diet-induced obesity. NMN-administered mice showed significantly higher energy expenditure during the dark period and a healthier plasma lipid profile, but without a reversal of glucose intolerance or insulin resistance. Importantly, the beneficial effects of NMN administration were strongly attenuated in mice that expressed catalytically-inactive SIRT1. Using a combination of plasma mass-spectrometry and O-link proteomics, we also identified key SIRT1 independent and dependent circulating proteins and pathways that were significantly altered by obesity and oral NMN administration. Glucose metabolism, cholesterol metabolism and immune related pathways are among the most significant changes. These data support the hypothesis that NAD^+^-dependent SIRT1 activity mitigates obesity and associated abnormalities in glucose and lipid metabolism.

## Methods

### Mouse experiments

All animal experiments were approved by the Institutional Animal Care and Use Committee (IACUC) at Weill Cornell Medicine – Qatar (Protocol ID. 2015-0026). The mouse colony was maintained in a facility accredited by the Association for Assessment and Accreditation of Laboratory Animal Care (AAALAC). Mice were housed at 23°C in plastic, single-use, individually ventilated cages (Innovive, USA) with unrestricted access to food and water, and maintained in a 12-h light, 12-h dark cycle (Lights on 0600 hrs.; lights off 1800 hrs.). Toes were clipped for identification and genotyping, which was carried out on DNA samples extracted using the HotSHOT method [45]. Tissue samples placed in a 96-well plate were digested for 30 minutes (@95°C) in 75 μL lysis buffer containing 25 mM NaOH and 0.2 mM EDTA, followed by the addition of 75 μL neutralization buffer (40 mM Trizma-HCl). Post-centrifugation, ∼25 μL DNA sample was carefully collected and stored at −20°C. To detect Cre-recombinase expression, the primer set used was: Cre Forward (oIMR1084): GCGGTCTGGCAGTAAAAACTATC; Cre Reverse (oIMR1085): GTGAAA CAGCATTGCTGTCACTT.

The internal positive control primer set was: Forward (oIMR7338): CTAGGCCACAGAATTGAAAGATCT; Reverse (oIMR7339): GTA GGTGGAAATTCTAGCATCATCC. A 96-well thermal cycler (Applied Biosystems) was used for PCR genotyping (20 μL reaction volume) and the thermal cycling parameters were as follows: <u>Step 1</u>: Denaturation (94°C, 1 minute); <u>Step 2</u> <u>(30 cycles)</u>: 94°C, 1 minute; 60°C, 1 minute; 72°C, 1 minute. <u>Step 3</u>: 72°C, 10 minutes. The PCR sample was mixed with 4 μL of the 6X loading dye and ∼20 μL was loaded on a 1% agarose gel containing 1 μg/ml ethidium bromide. Gel electrophoresis was performed for 40 minutes at 140V and PCR amplicons were detected on a ChemiDoc^TM^ MP Imaging system (Bio-Rad).

### Experimental details and NMN supplementation

A tamoxifen-inducible (Cre-ERT2) mouse model for whole-body deletion of exon 4 of SIRT1 has been described previously [46]. These mice express a catalytically-inactive SIRT1 protein. For experiments, floxed/Cre-ERT2 (Cre-positive) male mice were bred with floxed female mice (Cre-negative) to produce Cre-positive and Cre-negative littermates. Only male mice were used for experiments. SIRT1-deletion was achieved by feeding 8-week-old Cre-positive male mice a diet containing 360 mg/kg tamoxifen citrate for 5 weeks (Modified AIN-93G purified rodent diet, Dyets Inc.). Cre-negative mice were also fed tamoxifen to control for any potential side-effects of tamoxifen administration. To trigger obesity, 13-week-old mice were fed a high-fat diet (HFD) consisting of 60 Kcal% fat (Research Diets, Cat. D12492) or Chow (Pico-Vac Lab Rodent Diet, Cat. 5061) for 20 weeks prior to sacrifice. For the NMN project, NMN (400 mg/kg/day) was provided in drinking water concurrent with the HFD (GeneHarbor, Hong Kong). NMN was dissolved in Aquavive^®^ laboratory-grade ultrapure acidified water (pH 2.5-3.0), and the pH of NMN-containing water was not different from that without NMN. Fresh NMN-containing water was provided every 3 days and mice were weighed regularly to monitor changes in body weight and adjust NMN dosage. Mice were euthanized using CO_2_ asphyxia and dissected using sterile instruments to isolate tissues, which were immediately snap-frozen in liquid N_2_ and stored at −80°C until analysis.

### Intraperitoneal glucose-tolerance (IPGTT) and insulin tolerance tests (IPITT)

For IPGTT, mice were fasted for 16 hours overnight prior to performing GTTs (1700 – 0900 hrs.). A 2% glucose solution was prepared in phosphate-buffered saline (PBS) and sterilized by filtration. After measuring their fasting glucose levels, mice were injected with a dose of 2g/kg glucose and blood glucose levels were monitored 15-, 30-, 60-, and 90-minutes post-injection. Blood was sampled from the tail and measurements were made using an AlphaTRAK2 glucose monitoring kit.

For IPITT, mice were fasted for 6 hours prior to performing ITTs (0900 – 1300 hrs.). A 0.1 unit/ml solution of insulin was prepared by adding 16.6 μL of 10 mg/ml recombinant human insulin (Cat. I9278, Sigma-Aldrich) to sterile PBS. After measuring their fasting glucose levels, mice were injected with an insulin dose of 0.75 units/kg and blood glucose levels were monitored 15-, 30-, and 60-minutes post-injection. Measurements were performed as described above for IPGTT.

### Assessment of energy expenditure using the Comprehensive Lab Animal Monitoring System (CLAMS)

The Comprehensive Lab Animal Monitoring System (CLAMS) (Columbus Instruments, USA) is a rodent home-cage system that quantifies O_2_ intake (VO_2_) and CO_2_ production (VCO_2_), energy expenditure (EE), and respiratory exchange ratio (RER) in experimental animals by indirect calorimetry. RER is the ratio between VCO_2_ and VO_2_ (VCO_2_/VO_2_) and is a surrogate of substrate utilization. A value of 0.7 indicates reliance on fatty acid metabolism, while a value of 1.0 indicates dependence on carbohydrate metabolism. Before each experiment, the instrument was calibrated using a Class I calibration gas mixture containing 0.5% CO_2_ and 20.5% O_2_ (Buzwair Scientific and Technical Gases). The experiments were carried out over a 72-hour period; the first 24 hours were used to acclimatize mice to the cages [47] and the next 48 hours were used for experimental measurements. The CLAMS enclosure temperature was set to 23°C and all experiments were started at ∼0900 hrs. Food intake and physical activity were recorded using the in-built functions and animals had access to a running wheel for voluntary exercise. Water intake was assessed by weighing the water bottles before and after the experiment. For post-experiment data processing, the comma separated value (.csv) files generated by the Oxymax software were directly uploaded to the CalR web application [48, 49] to generate hourly-averaged data for individual mice to compare EE and RER between experimental groups (e.g., with or without NMN) and for statistical analysis and presentation.

### Body composition analysis (BCA) using Nuclear Magnetic Resonance (NMR) technology

The body composition (fat mass, fluid, and lean mass) of experimental mice was assessed using the Bruker Minispec Analyzer (Bruker Optics, Inc.), which uses nuclear magnetic resonance (NMR) technology. Before each experiment, a ‘daily check’ was performed using a standard provided by the manufacturer to validate the equipment functionality. Mice were transferred to a clear, plastic cylinder capped at one end and then immobilized by inserting a plunger through the other end. The cylinder was then inserted into the sample chamber of the instrument for the duration of the scan, which lasted ∼2-3 minutes. The cylinder was thoroughly cleaned between measurements to remove any urine or feces excreted by the sampled mouse.

### Plasma preparation

Mice were euthanized by CO_2_ asphyxiation and ∼0.5-1 ml blood was quickly collected by cardiac puncture and transferred gently to EDTA-coated blood collection tubes (Cat. 367844, BD Vacutainer) to which 100 μL of 2% EDTA had been previously added. A 3-ml syringe and 23G needle were used for cardiac puncture. Blood samples were centrifuged at 1200 rpm for 15 minutes at 4°C and ∼100-150 μL plasma was carefully collected, aliquoted, and stored at −80°C until analysis.

### Plasma O-Link proteomics

Protein measurements were performed on plasma samples using the mouse exploratory panel, developed by O-link Proteomics (Uppsala, Sweden) and based on the Proximity Extension Assay (PEA), which employs a pair of oligonucleotide-labeled antibodies specific to each target protein. A unique protein-specific sequence is generated by a DNA polymerization event when 2 matching oligonucleotides are in proximity, which is then quantified by quantitative PCR. The mouse exploratory panel allows the simultaneous quantification of 92 proteins at a time using as little as 1 μL sample volume. The Olink assay was performed at Weill Cornell Medicine-Qatar using the Mouse Exploratory Panel (Product # 95380) to measure protein biomarkers in randomized mouse plasma (EGTA) samples. The assay was carried out according to manufacturer protocol. The O-link assay was performed on a 96-well integrated fluidic circuits chip (Fluidigm, San Francisco, CA) and signal quantification was carried out on a Biomark HD system (Fluidigm, San Francisco, CA). Each sample was spiked with quality controls to monitor the incubation, extension, and detection steps of the assay. Additionally, samples representing negative controls and inter-plate controls were included in each analysis run. From raw data, real time PCR cycle threshold (Ct) values were extracted using the Fluidigm RT-PCR analysis software at a quality threshold of 0.5 and linear baseline correction. Ct values were further processed using the O-link NPX manager software. Here, Ct values of each sample and analyte were normalized based on spiked-in extension controls. Normalization is performed to minimize intra- and inter-assay variation. Inter-plate variability was calibrated using Ct values of inter-plate control samples. Finally, obtained values were inverted to obtain normalized log2 scaled protein expression (NPX) values. Therefore, a difference of 1 NPX between samples corresponds to a doubling of protein concentration.

### Plasma mass-spectrometry experiments and data analysis

Pierce BCA Protein Assay (Thermo Fisher Scientific, 23225, USA) was used to determine the plasma protein concentration. 50 μg protein was digested using iST sample preparation PreOmics kit 96x (PreOmics GmbH, Germany, P.O.00027). 50 μl of the lysis buffer was added to the plasma sample and heated at 95°C in a thermomixer (1000 rpm, 10 minutes). For protein digestion, 50 μl of trypsin/LysC mix was added at 37°C in thermomixer (500 rpm, 3 hours). To stop the reaction, 100 μl of stop buffer was used followed by washing steps to clarify the peptides and remove any hydrophilic or hydrophobic contaminants. Then, peptides were eluted using the elution buffer. After speed vac, purified peptides were resuspended in 50 μl of LC-load buffer at a final concentration of 1 μg/μl and iRT peptides were added. The peptide samples were randomized and analyzed by LC-MS/MS in two batches using the Q Exactive HFX mass spectrometer (Thermo Scientific) equipped with a nano-flow chromatographic system (Easy nLC-II 1200, Thermo Scientific). Peptides were ionized by 2.5 kV spray voltage and introduced into the mass spectrometer. For fragmentation of precursor ions, data-dependent analysis (DDA) was conducted following 15 most-intense precursor ions that were detected in the full scan. Dynamic exclusion was applied for 60 seconds, and then the selected ion was fragmented by higher energy collisional dissociation (HCD, collision energy 28).

Data were analyzed using Genedata Expressionist (v.13.0.1) and Mascot (v2.6.2) software. The raw MS data were processed using two Genedata modules: Refiner MS for pre-processing the data and Analyst for post-processing the data and basic statistical analysis. After noise reduction, LC-MS1 peaks were detected and their properties were calculated (m/z and RT limits, m/z and RT averages, intensity). Chromatograms were further aligned based on the RT spectra. Individual peaks were grouped into clusters, and MS/MS data assigned to these clusters were annotated with a Mascot search for MS/MS ions using a peptide tolerance of 10.0 ppm, an MS/MS tolerance of 0.10 Da, a maximum number of missed cleavages of 2, and the Uniprot database for mice. Results were validated by applying a threshold of 1% corrected normalized false discovery rate (FDR). Protein interference was performed based on peptide and protein annotations. Redundant proteins were ignored using Occam’s razor principle, and at least one unique peptide was required for positive protein identification. Protein intensities were calculated using the Hi3 method. The samples were further grouped, and the corresponding data were post-processed using Genedata Analyst. As a first step, the two batches were independently normalized based on the Reference Rows option using the Arithmetic Mean Averaging method and the Relative Function Divide (sum of the total list divided by the number of elements in the list). The individual corrected batches were then merged, and central tendency normalized using the median. To compare the different groups, T-tests were performed using the means and (at least 50%valid protein values per group) to calculate the effect size and corresponding p-values. The calculated effect size reflects the fold change of each protein.

### Sample preparation for mass spectrometric analysis

Plasma samples underwent trypsin digestion using the PreOmics iST kit (PreOmics GmbH, Martinsried, Germany) [50, 51]. Briefly, each sample was treated with LYSE buffer at 95°C for 10 minutes for alkylation and reduction of cysteine, and then treated with DIGEST buffer for trypsin digestion at 37°C for 3 hours. Peptides were then quenched by STOP buffer and washed with WASH buffer 1 and 2. In the final step, peptides were eluted by ELUTE buffer and dried using a speed vacuum chamber. Each dried peptide sample was reconstituted and sonicated in LC-LOAD buffer for LC-MS/MS.

### Mass spectrometric acquisition using data-independent acquisition (DIA) and data-dependent acquisition (DDA) and data processing

Mass spectrometric acquisition was performed by a nano-flow chromatographic system (Vanquish Neo UHPLC system, Thermo ScientificTM) coupled to Orbitrap Exploris 480 mass spectrometer (Thermo ScientificTM). One μg of each peptide mixture sample was injected onto EASY-spray PepMap Neo UHPLC C18 column (particle size 2 μm, 100Å, I.D.75 μm, length 50 cm, Thermo ScientificTM) in 45°C column temperature for separation by 120 min gradient and then introduced into mass spectrometer by EASY-Spray™ Source (Thermo ScientificTM) with 2.3 kV spray voltage. For DIA analysis, the full scan (MS1) and MS2 scan were conducted between 350 – 1200 m/z, and 120k resolution for MS1 and 30k resolution for MS2 were applied. For MS 1 scan, normalized automatic gain control (AGC) target 300%, 1 micro scans, and auto maximum injection time mode were used. For MS2 scan, isolation window 23.4 m/z, 27 % HCD collision energy, 1000% normalized AGC target, 1 microscans were used. For DDA analysis, the full scan was conducted between 400-1650 m/z at 120k resolution, standard AGC target and auto maximum injection time. For MS2 analysis, 30k resolution, 60 second dynamic exclusion, 2 isolation window (m/z), custom AGC target mode, 250% normalized AGC target, auto maximum injection time mode, 1 microscans were applied. For fragmentation of the precursor ions, 28% HCD collision energy was used. For data analysis, Spectronaut version 17 (Biognosys) software was used. DIA raw data were processed with DDA raw data by a spectrum centric, also known as library-free approach (direct DIA) using the Mus musculus UniprotKB fasta database (release 2023_08, 55,221 protein entries). The length of peptides was set to range 5 to 52 amino acids, with carbamidomethylation (+57.021 Da) of cysteine as a fixed modification, and acetylation (+42.011 Da) of N-terminal and oxidation (+15.995 Da) of methionine as valuable modification. Twenty ppm tolerance for MS1 and MS2 search, run wise imputing for imputation strategy, and sum peptide quantity for major group quantity were applied. In the post analysis step, t-tests and PCA analysis was calculated using the default parameters in Spectronaut and resulting fold changes and p-values were used for further analysis.

### Identifying significant differentially expressed proteins (SDE-proteins)

We define SDE proteins between two conditions as those that have a Benjamini-Hochberg adjusted p-value < 0.05, have more than one mapped peptide (for Mass-spec data) and have an absolute log2 FC >= 0.58 (i.e. at least a 50% change in any direction). For string-db analysis, ingenuity pathway analysis and grouped analysis, a threshold of p-value < 0.05 was used.

### String-db Pathway enrichment analysis

Enrichment analysis was performed for both individual and grouped comparisons using the online interface of string-db [52] (https://string-db.org, version 12.0). For individual comparisons (i.e. CH vs CC), all SDE proteins were submitted using the “Proteins with Values/Ranks” interface and analyzed with normal gene analysis parameters with the default whole-genome background applied. The default string-db moderate evidence setting was used for all interactions and network structure.

String-db networks are shown where each node represents one protein. Edge width represents the strength of evidence associating two proteins and includes physical interaction, co-expression and literature association. For paired comparisons, node center color represents the pathway membership color while node halo color represents the expression of the gene. For grouped analysis, node color represents the cluster membership. KEGG pathway and PFAM domain enrichment was also performed using string-db using an FDR < 0.05 as the pathway significance threshold.

### Grouped analysis and MCL Clustering

To increase power to detect expression trends and pathway enrichment, we conducted a grouped analysis. A protein in this analysis was included if it is significant (as previously described) in any of these four comparisons: CH vs CC, CH+N vs CH, SH vs SC, SH+N vs SH. This gene list was uploaded using the “Multiple proteins” interface to string-db. KEGG and PFAM enrichment was performed as previously mentioned. Markov Cluster Algorithm (MCL) was performed within string-db with an inflation parameter of 1.5. MCL recover clusters of genes within the string-db network using a simulation of stochastic flow [53]. In MCL analysis, a gene can belong to at most one cluster.

### Ingenuity Pathway Analysis (IPA) Causal Analysis and String-db Functional Annotation

IPA software (Qiagen) was used for upstream causal prediction function. Upstream analysis identifies potential master regulators that can explain the observed changes in our data based on prior knowledge of expected effects between gene regulators and their target genes stored in the Ingenuity® Knowledge Base. For each potential upstream master regulator two statistical measures are computed: an overlap p-value and an activation z-score. The overlap p-value calls likely upstream regulators based on significant overlap between dataset genes and known targets regulated by an upstream regulator. The activation z-score is used to infer likely activation states of master regulators based on comparison with a model that assigns random regulation directions. Core analysis was performed using the thresholds described for SDE proteins followed by upstream analysis filtering for only Gene products. We chose the union of the top 20 by z-score and top 20 by p-value for visualization and further analysis.

Predicted master regulators and the targets in our dataset were then further annotated. The molecule type was annotated with IPA software. Functional annotations were obtained from string-db and analyzed by matching (case insensitive search) any of several key terms (shown in parenthesis) for six functional categories: Immune function (’Immune’,’Immunity’,’Inflammation’,’T cell’,’B cell’,’Inflammatory’,’leukocyte’,’interleukin’), glucose function (’Glucose’,’Gluocogenesis’,’Sugar’), lipid function (’Adipose’,’Fat’,’Lipid’), energy expenditure (’Energy Expenditure’,’Energy’,’Expenditure’), metabolism (’Metabolism’,’Metabolic’) and Aging (’Aging’,’Senescence’,’Age’). Note that in the functional analysis the predicted master regulator CYP2C19 was replaced with the closest mouse homologue CYP2C29. Matches for each gene were summed to obtain a functional annotation score.

### Plasma lipid profiling

Plasma cholesterol, HDL, LDL, and triglycerides levels were quantified by the Laboratory of Comparative Pathology at Memorial Sloan Kettering Cancer Center (MSKCC, New York) using a Beckman Coulter AU680 clinical chemistry analyzer. The lipid profiling reagents used were OSR6116 (cholesterol), OSR60118 (triglycerides), OSR6195 (HDL), and OSR6196 (LDL). The technicians were blinded to the experimental details and genotype and were only provided a mouse identification number and an abbreviated experimental group label.

### Quantitative PCR

Post-euthanasia, tissue samples were snap-frozen in liquid nitrogen and stored at −80°C. For RNA extraction from adipose tissue samples, Qiazol lysis reagent (∼500 μL) was added to the samples prior to a rapid homogenization step (lasting ∼5 seconds). Samples were then centrifuged at 8500g for 15 minutes at 4°C, clarified by careful removal of the fat cake, and then transferred to PCR-clean 1.5 ml Eppendorf tubes without dislodging the cell debris pellet. RNA extraction, cDNA synthesis, and qPCR were performed as described previously [24]. Briefly, total RNA was isolated using the miRNeasy kit (Cat. 217004, Qiagen) following the manufacturer’s instructions, and a DNA digestion step was incorporated by incubating the sample with DNase I (Cat.79254, Qiagen) (RT, 15 minutes) prior to RNA elution. 500-1000 ng RNA was reverse-transcribed to cDNA using the High-Capacity RNA-to-cDNA kit (Cat. 4387406, ThermoFisher Scientific). QuantStudio6 Flex Real-Time PCR System (ThermoFisher Scientific) was used for qPCR and the thermal cycling parameters were as follows: <u>Stage I</u>: Denaturation (50°C, 20 seconds; 95°C, 10 minutes); <u>Stage II (40 cycles)</u>: 95°C, 15 seconds; 60°C, 1 minute; <u>Stage III (Melt curve analysis)</u>: 95°C, 15 seconds; 60°C, 1 minute; 95°C, 30; seconds; 60°C, 15 seconds. Sequences of all the primers used in the study are listed in Supplementary Table 1. qPCR data were quantified using the 2^-DCt^ method and, for clarity, 2^-DCt^ values x1000 are reported in the manuscript. GAPDH was used as the ‘housekeeping’ reference gene throughout.

### Experimental design, data analysis and statistics

Floxed Cre-ERT2 male mice were bred with floxed females to generate Control and SIRT1 iKO mice in the same litter (post-tamoxifen). Age-matched male mice were used for experiments and they were randomly assigned to experimental groups (Chow, HFD, NMN etc.), which were set up to run in parallel. The NMN data described in the manuscript were generated from 2 cohorts. In mice from cohort 1, we performed IPGTT, IPITT, BCA, plasma O-link and mass-spectrometry proteomics, and lipid profiling. After procuring the CLAMS, we repeated the NMN experiment in cohort 2 for *in vivo* measurements of energy expenditure, RER, and physical activity. IPGTT, IPITT, and BCA were repeated in cohort 2 and the results were qualitatively similar to those from cohort 1. The effect of oral NMN on weight-gain during DIO was statistically significant in both cohorts. The investigator(s) was blinded to genotype when weighing the mice and performing metabolic studies, including IPGTT and IPITT. It was not possible to blind the investigator(s) to cages with HFD or NMN because these cages were marked with special husbandry (SPHUB) cards and NMN-containing water was routinely provided in amber-colored bottles to distinguish those cages from the rest. Mice were not excluded during the experiments unless they had to be euthanized due to health issues such as malocclusion, rectal prolapse, or bite wounds due to fighting. Mice were group-housed except during the CLAMS experiments. Experimental mice were randomized during the CLAMS experiments and because the laboratory used a 16-cage system, analysis of NMN-fed Control mice (n=8 per group) was followed by experiments in SIRT1 iKO mice (n=8 per group) for the DIO study. Chow-fed control and SIRT1 iKO mice (n=6 per group) were analysed together. For plasma lipid profiling experiments performed at MSKCC, the technicians were blinded to the experimental details and genotype, and were only provided a mouse identification number and an abbreviated experimental group label. Data were analyzed using GraphPad Prism 9.0 and are presented as mean ± SEM. For statistical analysis of experiments comparing 2 groups, an unpaired Student’s *t*-test was used. In experiments that included more than 2 groups, Analysis of Variance (ANOVA) followed by Tukey’s *post hoc* test was used. For all statistical analysis, a ‘p’ value of <0.05 was considered statistically significant and the number of mice in any experimental cohort was at least 5. The specific number of mice in each experiment is indicated in the figure legends.

## Results

A tamoxifen-inducible (Cre-ERT2) mouse model for whole-body deletion of exon 4 of SIRT1 has been described previously [46]. These mice express a catalytically-inactive SIRT1 protein (Figure 1a). For experiments, floxed/Cre-ERT2 (Cre-positive) mice were bred with floxed mice (Cre-negative) to produce Cre-positive and Cre-negative littermates (Supplementary Figure 1a,b). To induce SIRT1-deletion, 8-week old male mice were fed a diet containing 360 mg/kg tamoxifen citrate for 5 weeks. This diet led to weight-loss in both Cre-positive and Cre-negative mice, but this was not significantly different between the two groups (Cre-negative, 2.97±0.51g *vs* Cre-positive, 3.37±0.41g) (Supplementary Figure 1c). Mice were switched to a Chow diet at the end of the 5-week period and sacrificed a week later to evaluate SIRT1-deletion *in vivo*. Expression analysis of Cre-ERT2 and SIRT1 (using primers spanning exon 4 of SIRT1) in epididymal fat (eWAT) revealed that, as expected, Cre-ERT2 was robustly expressed in Cre-positive but not in Cre-negative mice. The data also indicated that – in contrast to tamoxifen-fed Cre-negative (Control) mice – feeding tamoxifen to Cre-ERT2-positive mice significantly reduced SIRT1 expression in eWAT (Supplementary Figure 1d). These mice are hereafter referred to as SIRT1 inducible-knockout (SIRT1 iKO) mice. Importantly, expression levels of other members of the Sirtuin family were not significantly altered between Control and SIRT1 iKO mice (Supplementary Figure 1e).

**Figure 1:**
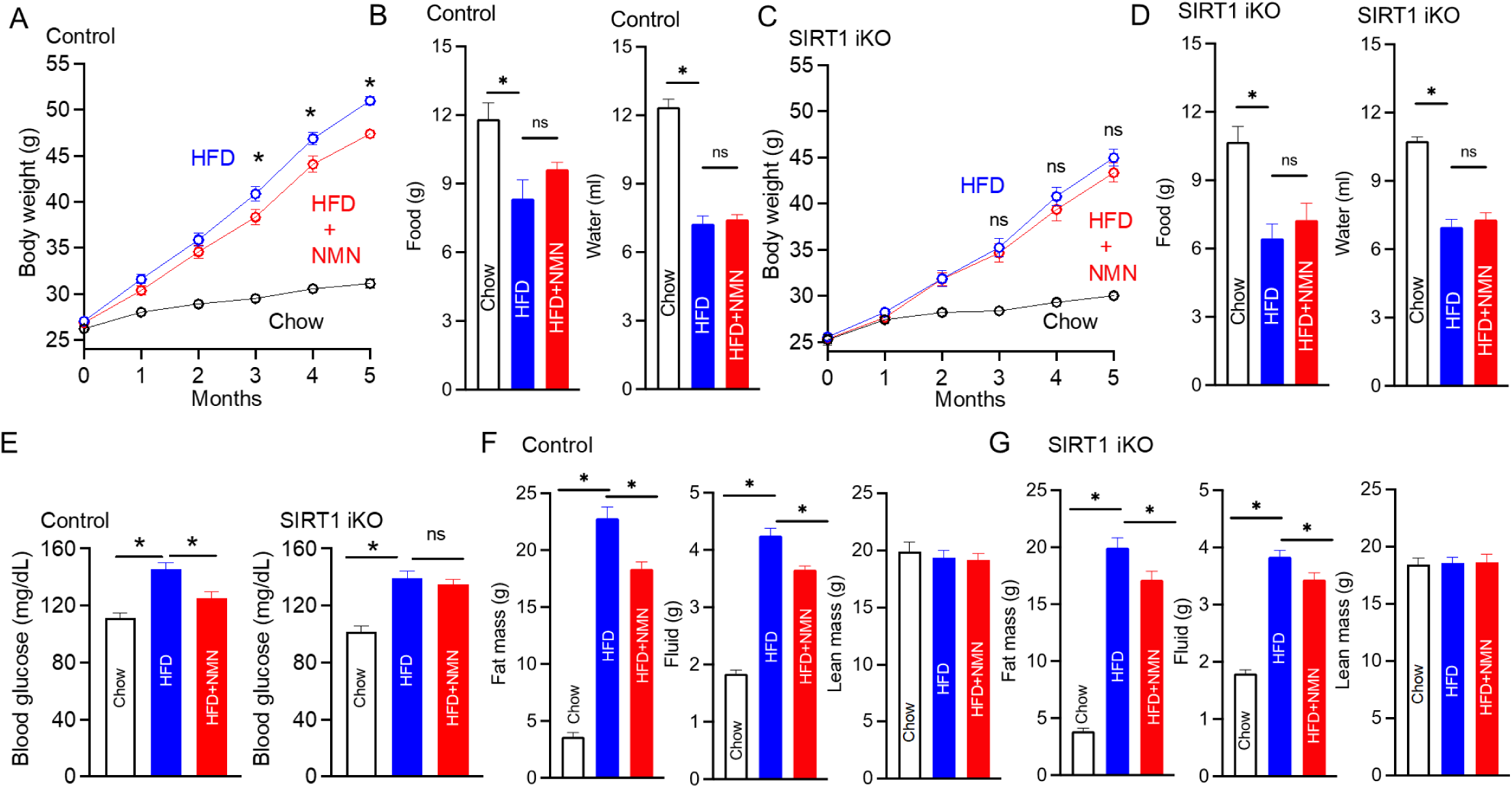
Oral NMN administration mitigates weight-gain during obesity in a SIRT1-dependent manner. (A) Data showing body weight measurements over the course of 5 months in Control mice fed chow (n=25), high-fat diet (HFD) (n=26), or HFD concurrent with 400 mg/kg/day NMN (n=25). (B) Food and water intake values recorded over a 3-day period in the indicated experimental groups (Chow, n=6; HFD, n=7; HFD+NMN, n=7). (C) Data showing body weight measurements over the course of 5 months in SIRT1 iKO mice fed Chow (n=16), high-fat diet (HFD) (n=21), or HFD+NMN (n=18). (D) Food and water intake values recorded over a 3-day period in the indicated experimental groups (Chow, n=6; HFD, n=8; HFD+NMN, n=8). (E) Blood glucose data collected from the indicated experimental groups after a 6-hour fast (Control mice: Chow – n=13, HFD – n=9, HFD+NMN, n=8; SIRT1 iKO mice: Chow – n=10, HFD – n=12, HFD+NMN, n=10). (F, G) Body composition analysis of Control and SIRT1 iKO mice (Control mice: Chow – n=14, HFD – n=20, HFD+NMN, n=17; SIRT1 iKO mice: Chow – n=13, HFD – n=24, HFD+NMN, n=17). Fat mass, fluid mass, and lean mass were estimated, as indicated.

### Oral NMN supplementation mitigates weight-gain in mice fed HFD in a SIRT1-dependent manner

A 20-week HFD study was performed in Control and SIRT1 iKO mice. A subset of mice receiving HFD were also provided the NAD^+^ precursor Nicotinamide Mononucleotide (NMN, 400 mg/kg/day) in drinking water to evaluate its effects on HFD-associated changes in weight-gain, energy expenditure, adipose tissue expansion, and glucose- and insulin-sensitivity. Control mice fed HFD gained a significant amount of weight over the course of 20 weeks and increased their body weights by ∼1.7-fold. However, Control mice that received HFD with NMN showed smaller body weight-gains and direct comparisons with mice fed HFD alone were statistically significant after 16 weeks (Figure 1A). Food and water intake were not different between the two groups (Figure 1B). Importantly, the suppressive effect of NMN on weight-gain was not observed in HFD-fed SIRT1 iKO mice, and body weights of SIRT1 iKO mice fed HFD alone were not significantly different from those fed HFD with NMN (Figure 1C). Food and water intake were not different between the two groups (Figure 1D).

Next, experiments were performed to investigate if the positive effect of NMN on weight-gain after HFD consumption was associated with improvements in whole-body glucose homeostasis. Blood glucose levels were monitored in the tail veins of fasted Control and SIRT1 iKO mice fed Chow, HFD, or HFD+NMN. In Control mice, consumption of HFD significantly increased fasting blood glucose levels. However, this increase was significantly smaller in Control mice that received HFD+NMN. SIRT1 iKO mice fed HFD also showed significantly higher fasting blood glucose levels but - in contrast to Control mice - the positive effect of NMN on fasting glucose levels was not observed (Figure 1E). In summary, the data suggested that weight gain and elevated fasting blood glucose levels observed in mice fed HFD were significantly suppressed upon administration of NMN in a SIRT1-dependent manner.

To evaluate obesity- and NMN-dependent effects on whole-body fat accumulation, body composition analysis was performed on Control and SIRT1 iKO mice fed Chow, HFD, or HFD+NMN. In Control mice, consumption of HFD led to a significant increase in fat mass. However – in agreement with the positive effects of NMN on body weight – mice fed HFD+NMN showed significantly less fat accumulation when compared to those fed HFD alone (Figure 1F). Importantly, parallel experiments in SIRT1 iKO mice indicated there was a significant difference in fat mass between SIRT1 iKO mice fed HFD+NMN and those fed HFD alone, although weaker than in Control mice (Figure 1G). We also observed a significant increase in fluid mass in HFD diet fed mice that was significantly reduced by NMN in both the Control and SIRT1 iKO mice (Figure 1F,G). Conversely, no significant effect on lean mass was observed in any condition (Figures 1F,G).

### SIRT1-dependent increase in energy expenditure in HFD-fed mice receiving oral NMN

Next, we assessed energy expenditure in Control and SIRT1 iKO mice fed HFD or HFD+NMN using the Comprehensive Lab Animal Monitoring System (CLAMS). Control mice that consumed HFD+NMN showed significantly higher energy expenditure when compared to those that consumed HFD alone (Figure 2A, B). Importantly, this increase was not observed in SIRT1 iKO mice (Figure 2C, D). Analysis of the respiratory exchange ratio (RER) values suggested that Control mice receiving HFD+NMN showed significantly higher RER when compared to those that received HFD alone (Figure 2E, F). Again, this difference in RER between the HFD and HFD+NMN groups was blunted in SIRT1 iKO mice (Figure 2G, H).

**Figure 2:**
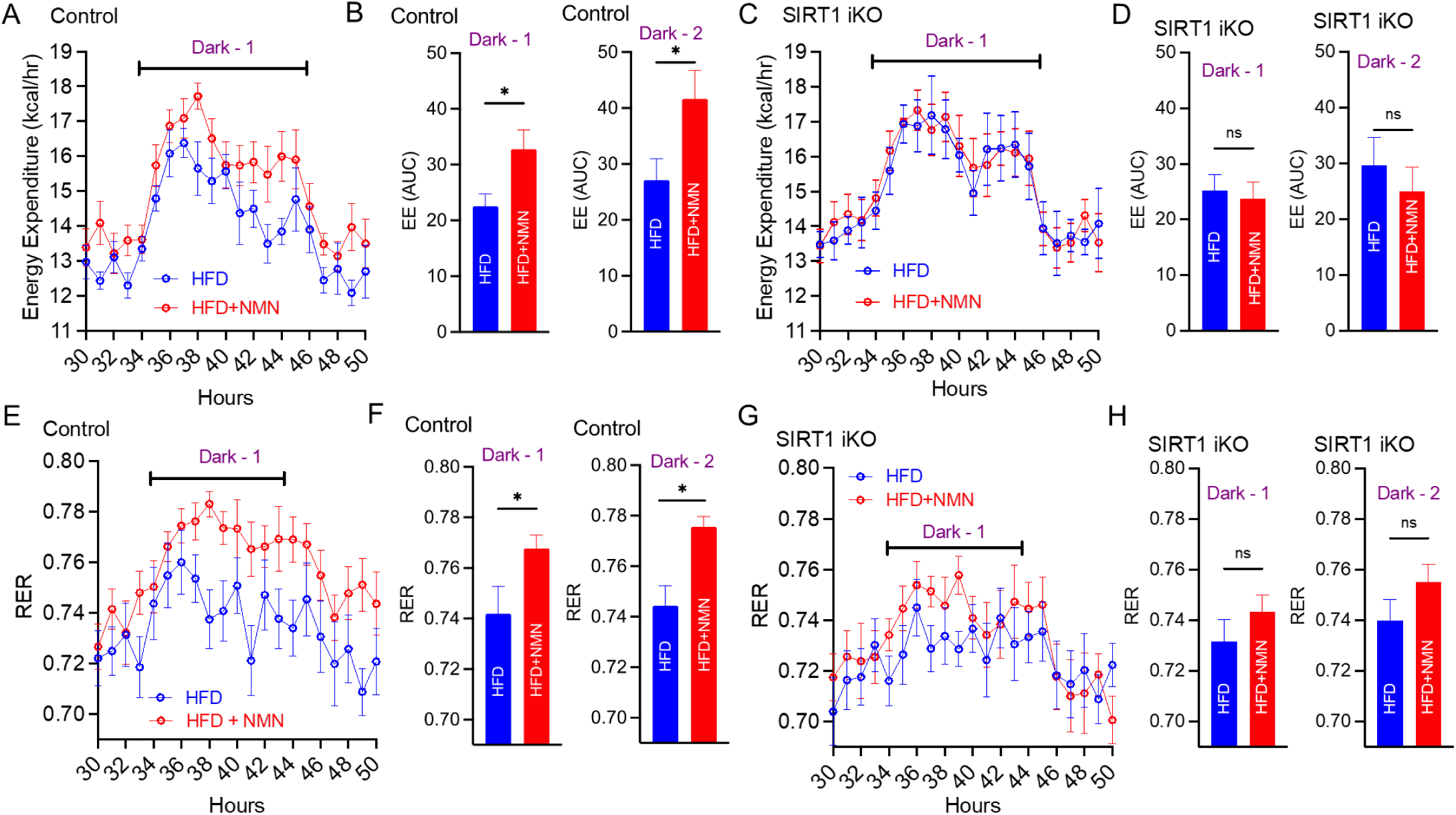
Oral NMN administration enhances energy expenditure in obese mice in a SIRT1-dependent manner. (A) Energy expenditure (kcal/hr.) in Control mice fed HFD (n=8) or HFD+NMN (n=8). (B) Area under curve (AUC) values for energy expenditure (EE) recorded during 2 consecutive dark periods for the experiments exemplified in (A). (C) Energy expenditure (kcal/hr.) in SIRT1 iKO mice fed HFD (n=8) or HFD+NMN (n=8). (D) Area under curve (AUC) values for energy expenditure (EE) recorded during 2 consecutive dark periods for the experiments exemplified in (C). (E, F) Respiratory exchange ratio (RER) values recorded during 2 consecutive dark periods in Control mice fed HFD (n=8) or HFD+NMN (n=8). (G, H) (F) Respiratory exchange ratio (RER) values recorded during 2 consecutive dark periods in SIRT1 iKO mice fed HFD (n=8) or HFD+NMN (n=8).

### Oral NMN supplementation did not improve glucose-tolerance or insulin-sensitivity but partially reversed dyslipidemia in obese mice

Mice were also subjected to glucose-tolerance (GTT) and insulin-sensitivity tests (ITT). In Control mice, consumption of HFD was associated with significant glucose-intolerance. However, administration of NMN did not show an overall improvement in glucose-tolerance and area under the curve (AUC) calculations indicated no significant differences between Control mice fed HFD and those fed HFD+NMN (Figure 3A). Parallel GTT studies performed in SIRT1 iKO mice also indicated similar results and NMN administration did not improve glucose-intolerance observed in HFD-fed SIRT1 iKO mice (Figure 3A). Whole-body insulin-sensitivity was compared by performing insulin-tolerance tests (ITT) in Control and SIRT1 iKO mice fed Chow, HFD, or HFD+NMN. Compared to Chow-fed mice, Control mice fed HFD showed blunted responses to insulin, which were most apparent at the early time-points of 15 and 30 minutes. However, the effect of insulin on blood glucose in Control mice fed HFD+NMN was not significantly different from mice fed HFD alone after 15 and 30 minutes (Figure 3B). Parallel ITT studies in SIRT1 iKO mice also indicated similar results and NMN administration did not produce any significant improvement in insulin-sensitivity in HFD-fed SIRT1 iKO mice (Figure 3B). Plasma lipid profiling revealed that Control but not SIRT1 iKO mice fed HFD+NMN had significantly lower circulating levels of cholesterol and LDL, although HDL levels were unaffected (Figure 3C, D).

**Figure 3:**
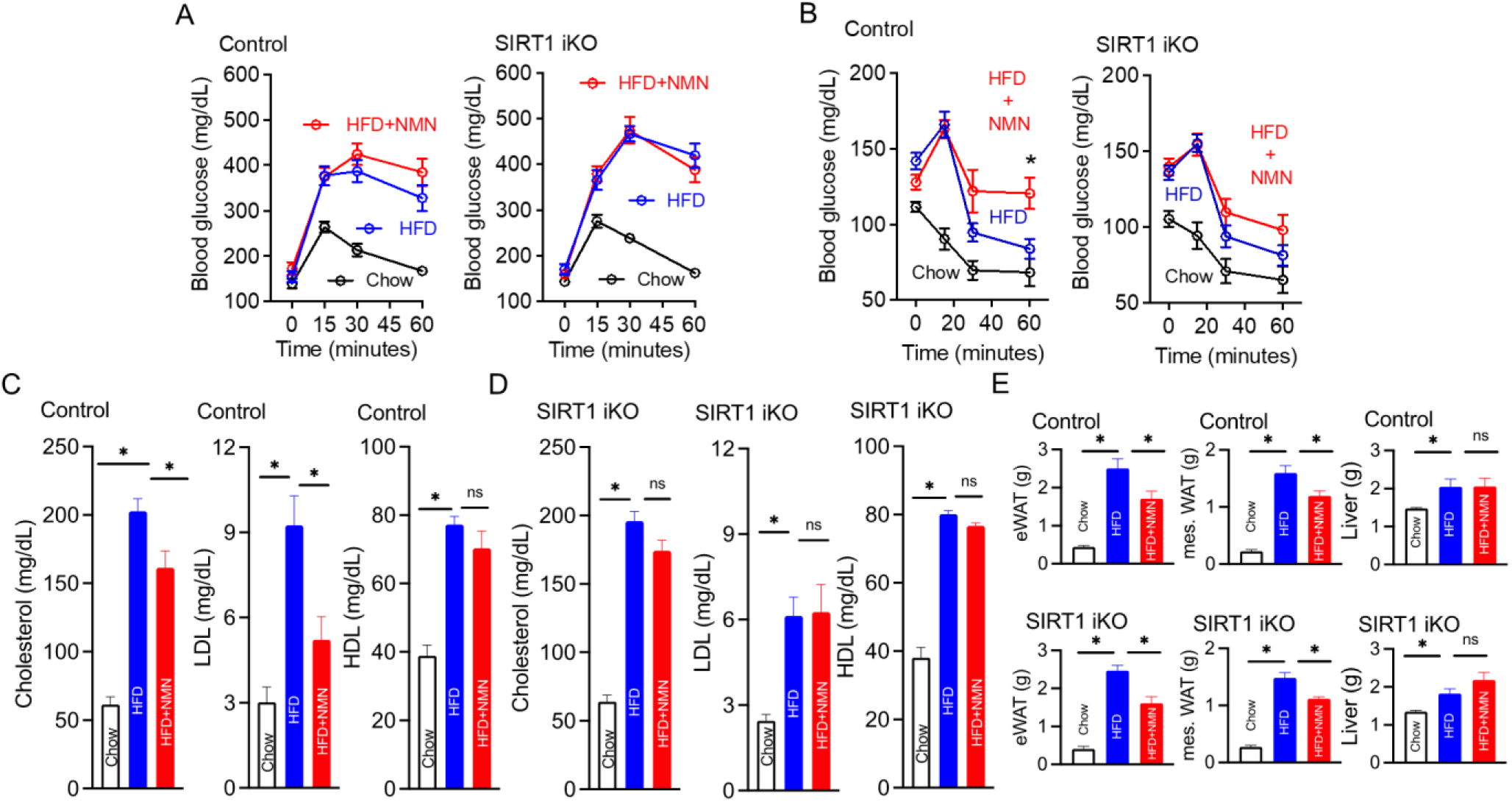
Oral NMN administration improves dyslipidemia but not glucose-intolerance or insulin-sensitivity. (A) Data from intraperitoneal glucose-tolerance tests (IPGTT) performed in Control and SIRT1 iKO mice fed Chow, HFD, or HFD with NMN. Mice were fasted overnight for 16 hours and injected with 2g/kg glucose (Control mice: Chow – n=13, HFD – n=9, HFD+NMN, n=8; SIRT1 iKO mice: Chow – n=10, HFD – n=12, HFD+NMN, n=10). (B) Data from intraperitoneal insulin-tolerance tests (ITT) performed in Control and SIRT1 iKO mice fed Chow, HFD, or HFD with NMN. Mice were fasted for 6 hours and injected with 0.75U/kg insulin (Control mice: Chow – n=13, HFD – n=9, HFD+NMN, n=8; SIRT1 iKO mice: Chow – n=10, HFD – n=12, HFD+NMN, n=10). (C, D) Lipid profile of Control and SIRT1 iKO mice fed Chow, HFD, or HFD+NMN (Control mice: Chow – n=6, HFD – n=6, HFD+NMN, n=6; SIRT1 iKO mice: Chow – n=6, HFD – n=6, HFD+NMN, n=6). (E) Epididymal fat (eWAT), mesenteric fat (mes. WAT), and liver weights of Control and SIRT1 iKO mice from the indicated experimental groups (Control mice: Chow – n=13, HFD – n=9, HFD+NMN, n=8; SIRT1 iKO mice: Chow – n=10, HFD – n=12, HFD+NMN, n=10).

### Adipose tissue expansion in Control and SIRT1 iKO mice fed HFD or HFD+NMN

The effect of obesity and NMN administration on individual fat depots (eWAT and mesenteric fat) was also evaluated. In Control mice, consumption of HFD led to significant expansion of eWAT and this increase was significantly reduced in mice that received HFD+NMN (Figure 3E). Feeding HFD to SIRT1 iKO mice also led to eWAT expansion and, interestingly, NMN administration significantly limited this expansion (Figure 3E). The effect of obesity and NMN administration on the mesenteric fat (mesWAT) depot was also assessed and these data indicated that, in Control mice, consumption of HFD led to significant expansion of mesWAT and this increase was significantly reduced in mice that received HFD+NMN (Figure 3E). Feeding HFD to SIRT1 iKO mice also led to mesWAT expansion and NMN administration significantly limited this expansion (Figure 3E).

To summarize, the parameters mWat, fluid mass, fat mass and eWat are all significantly changed in HFD+NMN and HFD+NMN in SIRT1 KO, hence we classify them as SIRT1 independent. On the other hand, SIRT1 dependent changes include total cholesterol, RER, LDL, HDL, fasting glucose, energy expenditure and body weight which are all significantly changed in HFD+NMN in controls but not in HFD+NMN in the SIRT1 KO background. Lastly, water intake, food intake, liver, lean mass and water and blood glucose IPG and IPT are not changed in either condition and hence are also categorized as SIRT1 independent.

### Significant changes in the global protein expression profile in plasma of HFD+NMN-fed mice

We conducted plasma proteomics on the experimental cohort using two platforms: O-Link, an aptamer-based technology for a panel of 92 mouse proteins and mass spectrometry as summarized in Figure 4A. The overall group responses were very similar for the two platforms as shown in the PCA plots in Figure 4B. Chow-fed mice in both the Control and SIRT1 iKO are in distinct clusters separate from the HFD-fed mice. In contrast, HFD+NMN-fed mice had relatively small but consistent effects on global protein expression profiles. All proteomics data was therefore combined and used for identifying significant differentially expressed proteins (SDE proteins) and functional enrichment analysis.

**Figure 4:**
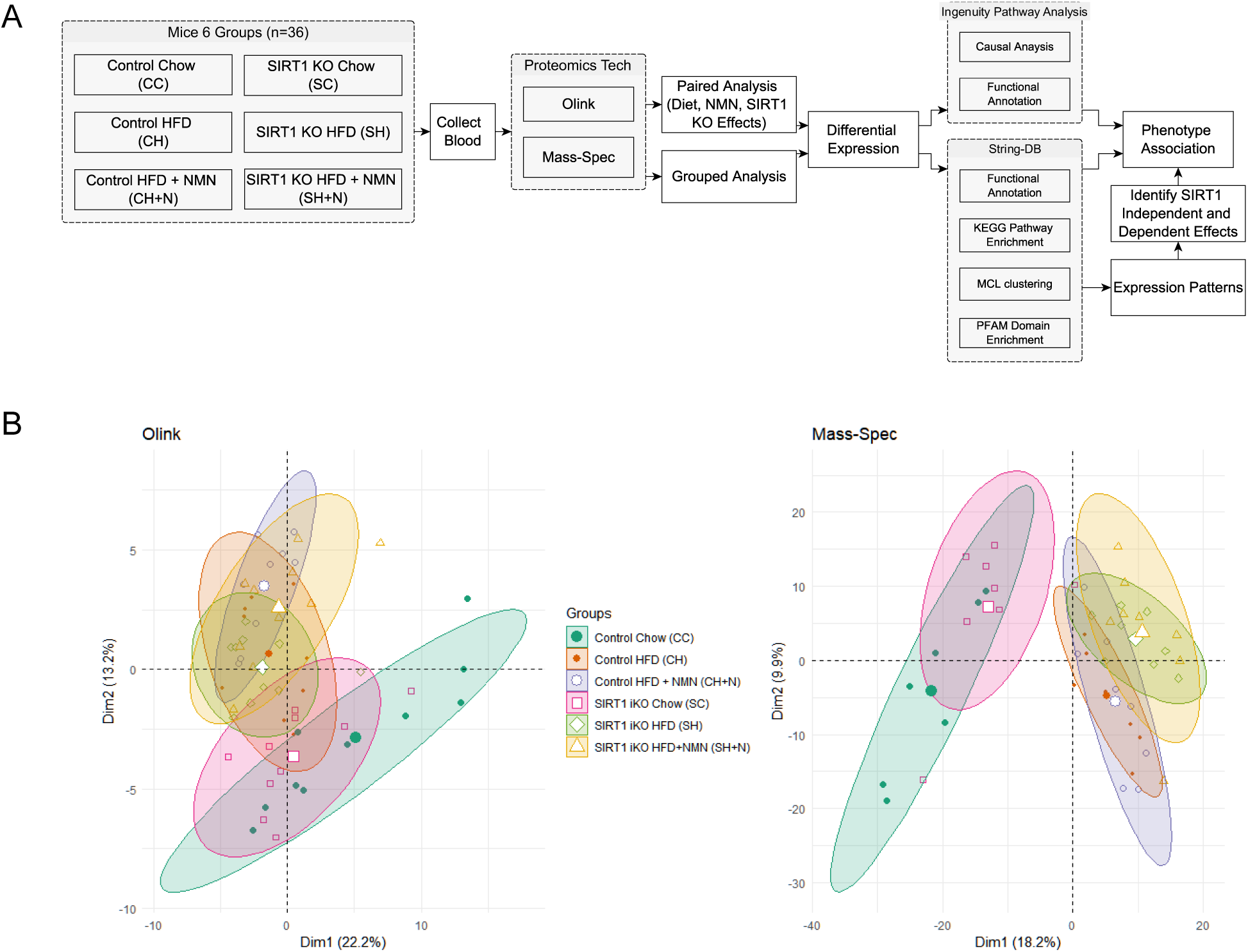
Proteomics Design and Sample PCA. A) Proteomics analysis flowchart depicting the main design and analysis steps B) PCA plots for Olink and Mass-Spec data. The first two principal components are shown. Ellipses are drawn to depict the shape of each group. The symbols and colors of each group are shown in the legend.

### High Fat Diet modulates linked pathways including Cholesterol metabolism, PPAR signaling proteins and Complement and Coagulation Cascade pathways in Control Mice

First, we examined the impact of HFD consumption on the plasma proteome of Control mice. As shown in the volcano plot, there were large expression changes due to HFD with 65 upregulated SDE proteins and 53 downregulated SDE proteins (Figure 5A). The top 10 non-immunoglobulin upregulated proteins are Creb3l3, Camp, Apoc2, Apoa5, Me1, Prg4, Saa1, Aldob, Lgals3 and Apoe while the top 10 non-immunoglobulin downregulated proteins are Serpina1e, Mup8, Tnc, Lifr, Mup2, Egfr, Serpina3k, Cfd, Cdh1 and Col3a1. Note the presence of multiple apoliporoteins among the top 10 upregulated proteins and two Mup proteins among the top 10 downregulated proteins. To determine if there are enriched pathways, we performed pathway enrichment analysis with STRING-DB. Among the top enriched KEGG pathways by pathway FDR include the complement and coagulation cascades, cholesterol metabolism and biosynthesis of amino acids (Figure 5B). Please note that proteins could belong to more than one KEGG pathway (Figure 5C). For example, Apoa1, Apoa2, Apoa3 are members of both the KEGG cholesterol metabolism and KEGG Pparg signaling pathways.

**Figure 5:**
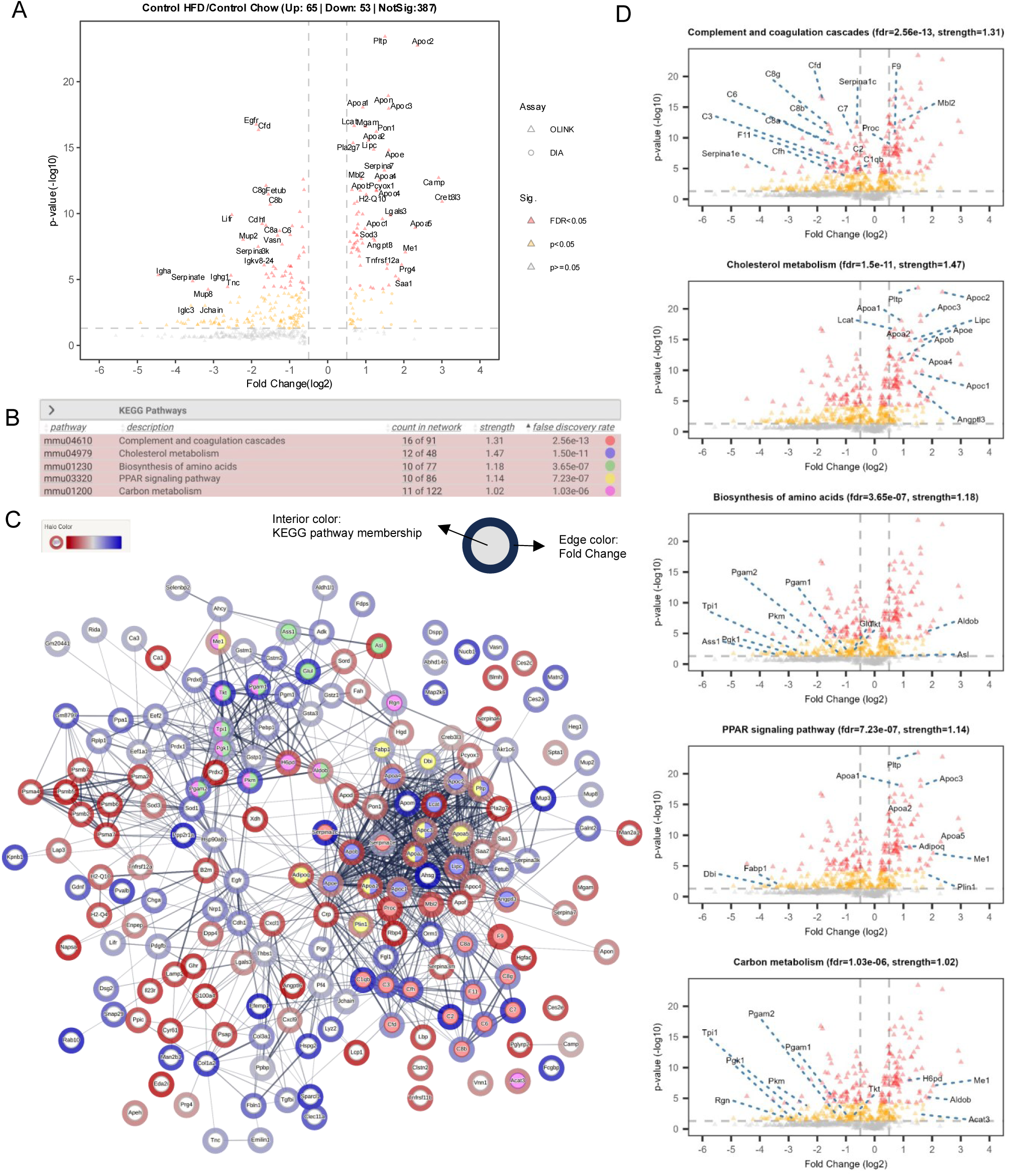
Diet effect in WT mice (Control HFD vs.Control Chow). A) Volcano plot showing the top 50 significant DE proteins. B) String-db network analysis for all high confidence significant differentially expressed proteins (peptides>1, p-value <0.05, abs(FC) > 0.5) between High-fat diet and Chow in wild-type mice. The top 5 enriched Kegg pathways by false discovery rate are shown. C) Network diagram showing the interactions between proteins and enriched network membership. Proteins belonging to a specific network are colored in the interior based on the color shown in B. Proteins may have multiple interior colors. The fold-change value is shown as a colored halo. Edge width is proportional to the strength of the evidence for an interaction. D) Volcano plots labelling only the members of the top 5 enriched kegg pathways. The legend is the same as in A.

To examine the general expression trends within pathways, we overlayed the expression of members of individual pathways onto the volcano plot. We see that while complement/coagulation members and amino acid biosynthesis genes are mostly decreased in response to HFD, cholesterol metabolism and PPAR signaling pathway proteins are all mostly increased (Figure 5D). One protein that is significantly upregulated in response to HFD but does not belong in enriched KEGG, reactome, wikipathway or PFAM is ApoN. ApoN is classified as a pseudogene in humans but is expressed in mice liver and is thought to be involved in cholesterol metabolism [54] .

Interestingly the network diagram shown in Figure 5C depicts numerous links between proteins belonging to separate KEGG pathways. For example, there are links between the completement and coagulation cascades proteins and cholesterol metabolism proteins. These links primarily reflect co-expression and co-mention in literature abstracts. Furthermore, many highly differentially expressed genes are not members of any of the top pathways depicted. Therefore, the overall complexity of interactions among the HFD induced proteins is not fully captured by looking at only the top pathways or at pathways individually. We later examine in an integrated method all gene expression patterns in a non-pathway centric method.

### Oral NMN supplementation significantly alters plasma levels of proteins linked to metabolic pathways

We next examined the effect of NMN treatment on HFD-fed Control mice. We found that NMN supplementation was associated with significant upregulation of 18 proteins and downregulation of 16 proteins (Figure 6A). The top non-immunoglobulin upregulated proteins are Cct7, Il17a, Serpina1e, Mup2, Mup8, Ca3, Mup20, Cct4, Capn2 and Slc2a3, while the top downregulated proteins are Ppbp, Gsr, Thbs1, Angpt1, Pf4, Pde5a, Mtpn, Pgm2, Eef1b2, Rida and Pdgfb, Spta1 and Fabp3. Note the presence of three MUP proteins among the top 10 upregulated proteins. The top five significantly enriched KEGG pathways are cysteine/methionine metabolism, ECM-receptor interactions, focal adhesion, glycine/serine/threonine metabolism and glycolysis/glucogenesis (Figure 6B). The string-db network is shown in Figure 6C and shows the interactions between different networks and proteins as well as whether a protein belongs to more than one network. As seen in the previous HFD analysis, there are numerous interactions between different proteins and pathways. The individual proteins belonging to the top five enriched KEGG pathways are shown in Figure 6D.

**Figure 6:**
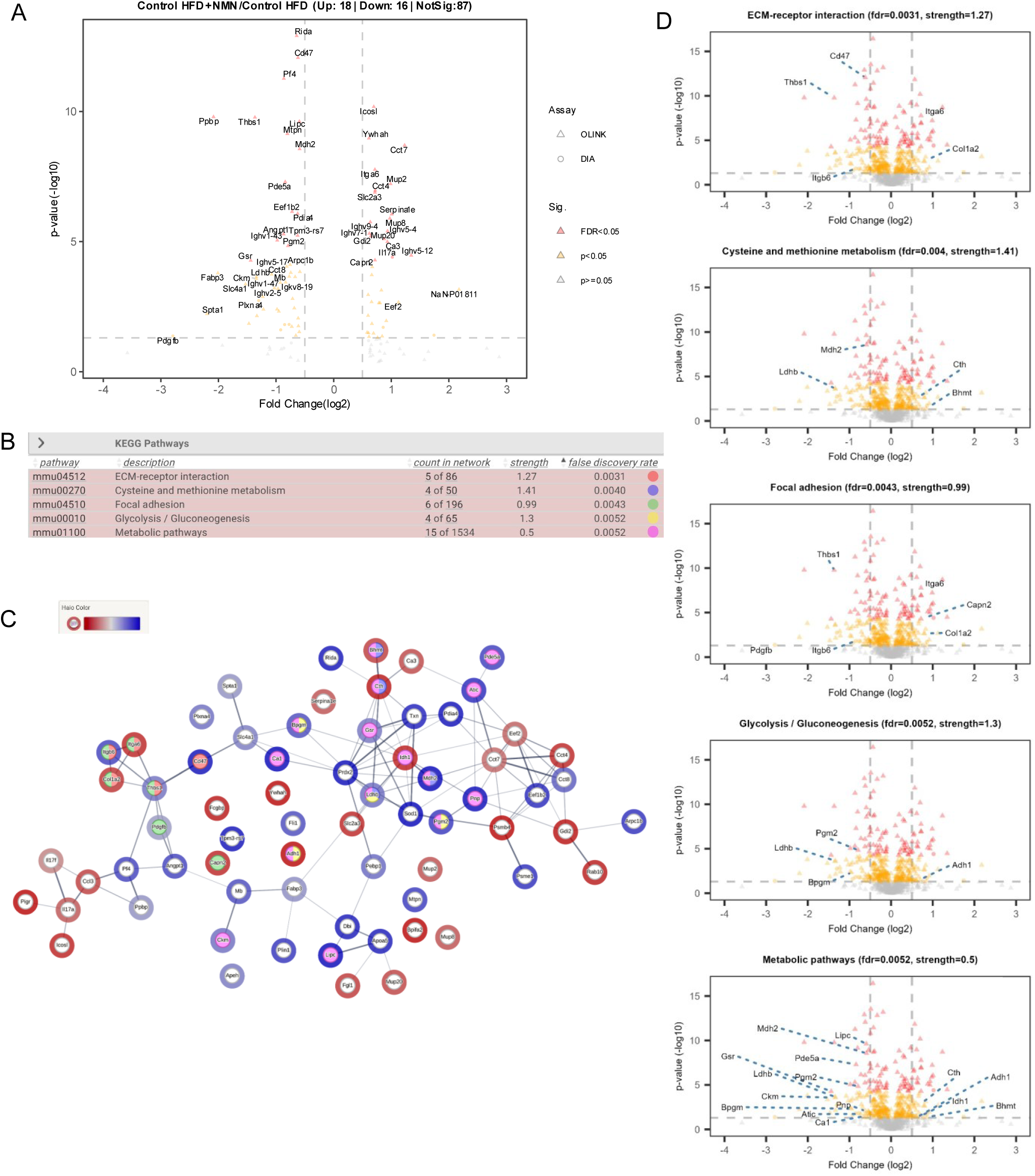
NMN effect in WT mice (Control HFD+NMN vs. Control HFD). A) Volcano plot showing the top 50 significant DE proteins. B) String-db network analysis for all high confidence significant differentially expressed proteins (peptides>1, p-value <0.05, abs(FC) > 0.5) between High-fat diet and Chow in wild-type mice. The top 5 enriched Kegg pathways by false discovery rate are shown. C) Network diagram showing the interactions between proteins and which proteins belong in which network. Proteins belonging to a specific network are colored in the center based on the color shown in A and B. Proteins may have different center colors. The fold-change value is shown as a colored halo. Edges are colored based on the type of interaction as shown in the legend. D) Volcano plots labelling only the members of the top 5 enriched kegg pathways. The legend is the same as in A.

### Plasma proteins linked to metabolic pathways downregulated in SIRT1 iKO mice

In parallel with the control mice, we also conducted proteome profiling experiments on the SIRT1 KO mice. SIRT1 knockout had a large downregulatory effect on protein expression on mice in all tested conditions (Figure 7). SIRT1 KO in the CHOW condition downregulated 23 proteins and upregulated 1 protein (Figure 7A). The top 10 downregulated proteins are Pygl, Fh, Cyb5a, Pvalb, Creb3l3, F11, Ctsd, Saa2, Mup4 and Dpp4 while 1 upregulated SDE protein is Serpina3m. Enrichment analysis of the SDE proteins demonstrate a metabolic phenotype with the top five enriched KEGG pathways being glucagon signaling, glycolysis/glucogenesis, carbon metabolism, glycine, serine and threonine metabolism and metabolic pathways (Figure 7B). Visualization of the network of interacting proteins (Figure 7C) shows numerous interactions between the SDE proteins.

**Figure 7:**
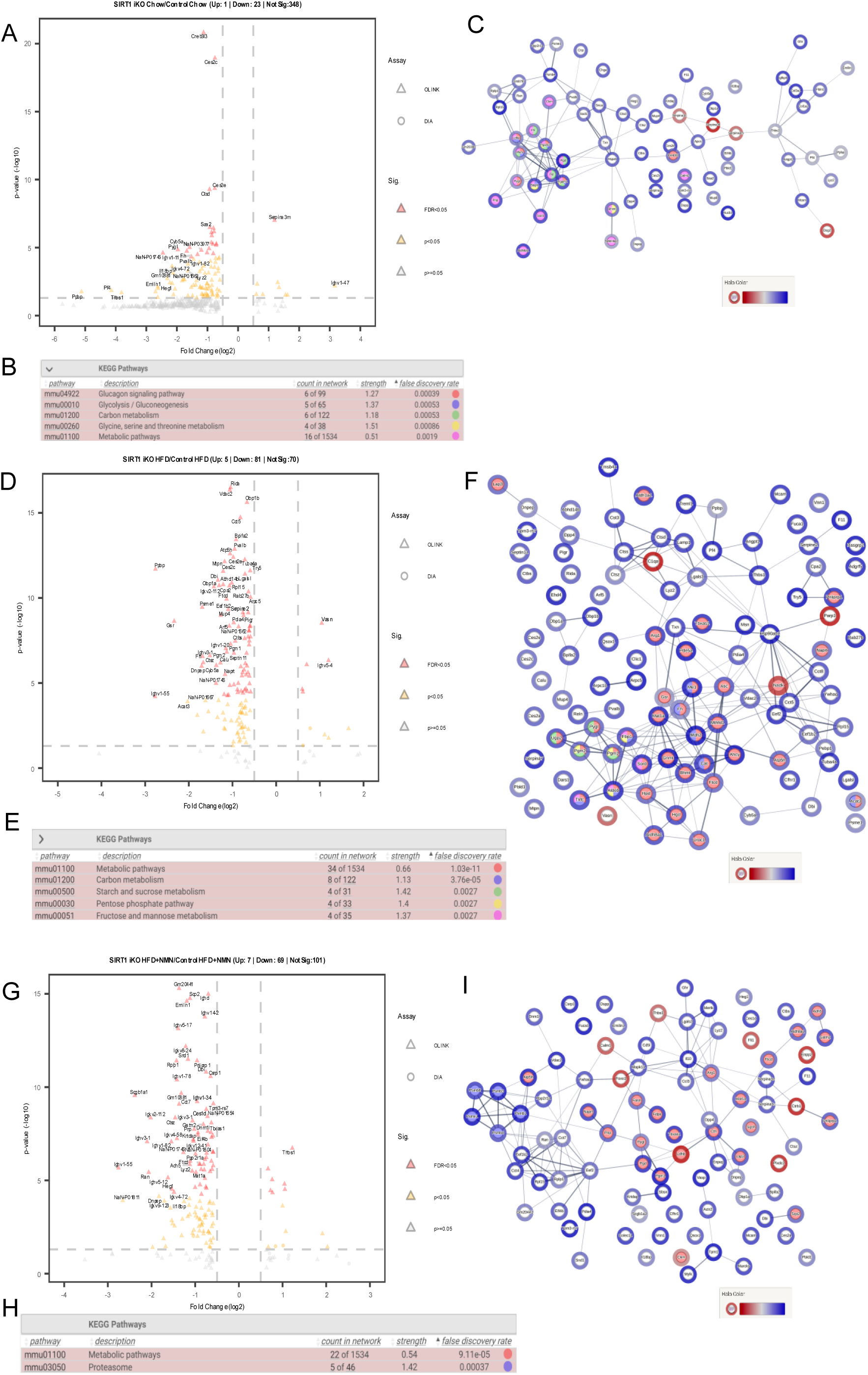
Comparison of SIRT1 KO effect. A,B) Chow, C,D) high fat diet and E,F) NMN-treat high fat diet. A,C, E) Volcano plots show the differential expression. The top 50 significant protein groups are labelled. Point shapes refer to the assay used (triangle=DIA, circle=OLINK). Point colors refer to the significance thresholds (red: fdr<0.05, orange: pvalue<0.05 and grey: pvalue>=0.05). Vertical dashed lines are set at an abs(FC)>0.5 and horizontal dashed line is set at-log10( p-value) < 0.05. Only groups with more than one peptide are included in this figure. B,D,F) The top 5 enriched Kegg pathways in the string-db network analysis.

Similarly, SIRT1 KO impacted mice fed a high fat diet with five upregulated proteins and 81 down regulated proteins(Figure 7D). The top 10 downregulated proteins are Ppbp, Gsr, Dnpep, Psme1, Fh, Ctsz, Cyb5a, Dbi, Obp1a and Calu while the top 2 non-immunoglobulin upregulated are Vasn and C1qa. As in the CHOW condition, SDE proteins were enriched in metabolic pathways with the top five enriched KEGG pathways being metabolic pathways, carbon metabolism, starch/sucrose metabolism, pentose phosphate pathway and fructose/mannose metabolism (Figure 7E). While many of the SDE proteins have known interactions, many also have no strong links to other proteins within the network (Figure 7F).

Finally, there is a large impact of SIRT1 KO on NMN supplementation in HFD. Notably, SIRT1 KO reduces the expression of many changes relative to WT with 7 upregulated proteins and 69 downregulated proteins (Figure 7G). The top seven non-immunoglobulin upregulated SDE genes are Thbs1, Calm1, Enpp2, Plxdc2, Ctrb1, Ldhb and the top 10 non-immunoglobulin downregulated SDE genes are Ran, Heg1,Ctsz, Rplp1, Gm20441, Cct7, Emilin1, Snd1, Adh5 and Scp2. Enrichment analysis on SDE proteins shows that the KEGG proteosome and metabolic pathways are enriched (Figure 7H). These SDE proteins also interact with each other across different pathways.

### Discovery of gene clusters linked to HFD using unsupervised hierarchical clustering

While the previous analysis focused on the highest individual protein changes and enriched pathways, there were numerous significant proteins that were not in the top enriched pathways. We therefore performed a grouped analysis combining significant results from different comparisons.

This resulted in a set of 344 genes to analyze. We also performed network-based clustering using the MCL algorithm as described in the methods (Figure 8). The results of MCL show that the network structure is composed of several subnetworks with two large networks (Figure 8A, 8B). Different clusters are enriched in different pathways as shown in Figure 8C. The top enriched KEGG pathways in MCL cluster 1 are metabolic pathways while the top enriched KEGG pathways in MCL cluster 2 are cholesterol and immune related pathways.

**Figure 8:**
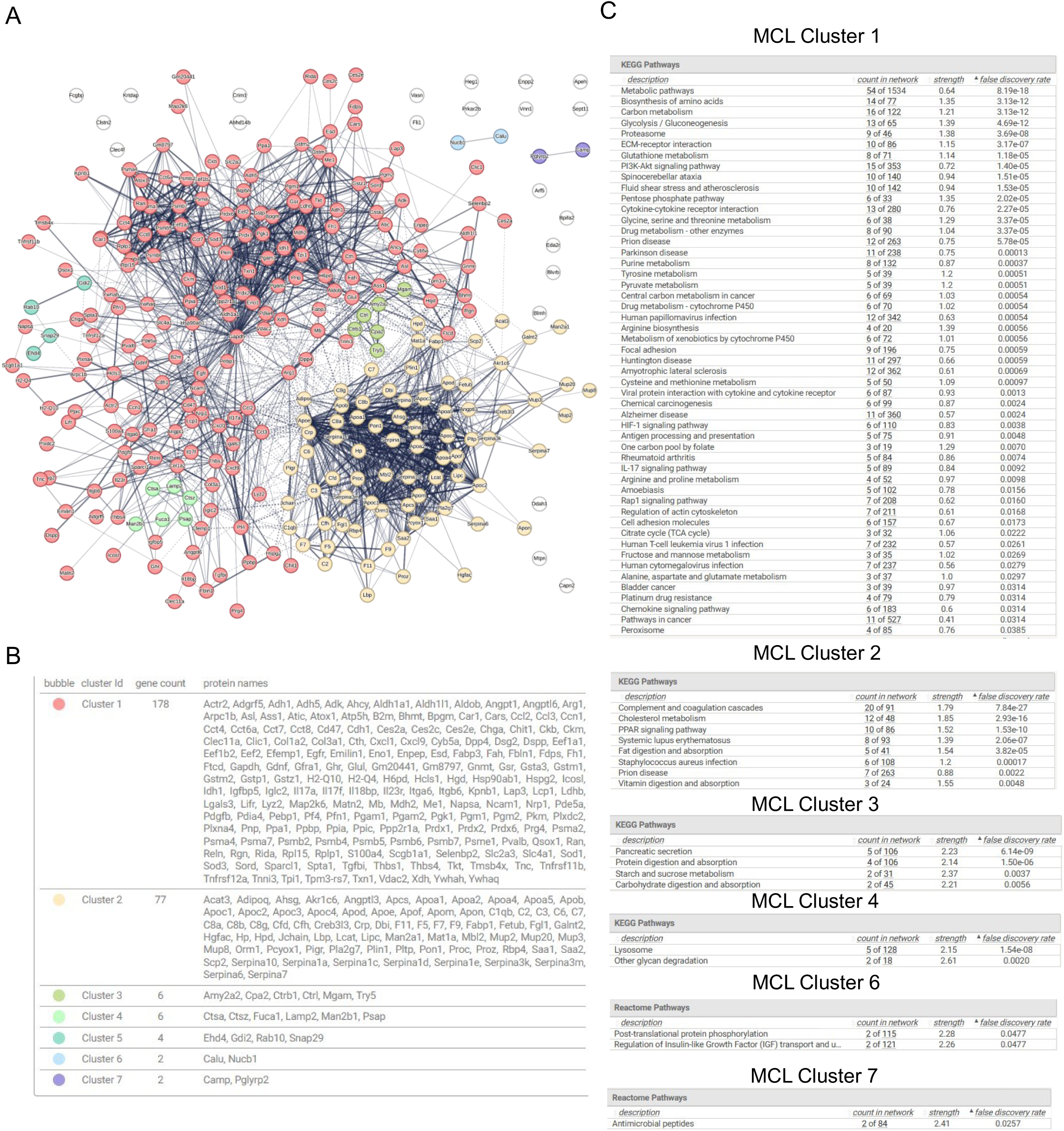
Grouped Analysis combining all significant proteins for the diet and NMN in control and chow. A) MCL clustering diagram showing the different clusters. Each cluster is depicted in a different cluster. Nodes without color have no interactions with other nodes at the medium confidence level. Solid line width between nodes is proportional to the evidence. Dashed lines depict interactions between groups. B) The list of clusters with gene counts and genes belonging to each cluster. C) Pathway enrichment for each MCL Cluster. Where possible, KEGG pathways are shown. If no KEGG pathways are enriched, then enriched reactome pathways are shown. Note there are no enriched functional pathways for MCL Cluster 5.

We also performed unsupervised hierarchical clustering on the expression values for each of the four comparisons annotating the MCL clusters and the proteomics technology use (supplementary Figure 2). The sample clustering clearly groups the samples with diet effect and NMN effects with an overall similar pattern for the SIRT1 KO and Control samples. The two main gene cluster groups are driven by genes that change expression due to the high fat diet. The top expressed changes with HFD is a gene cluster containing Creb3l3, Saa1, Saa2, Camp, Apoc2, Prg4 and Apoa5. These genes are mostly in MCL cluster 2 and are members of enriched pathways affecting PPAR signaling, cholesterol metabolism and PI3-Akt signaling. The bottom cluster are genes that are reduced in HFD and include Ppbp, Pdgfb, Try5, Gm20441, Jchain, Rida, Dbi, Ass1, Abhd14b, Gsta3, Gstm1 and Serpina1e. This includes genes that are members in several enriched immune, metabolic and signaling pathways and glutathione metabolism.

### SIRT1-independent and dependent changes in plasma proteins in mice fed HFD+NMN

Beyond the two main expression patterns are specific SIRT1-independent and SIRT1-dependent protein expression patterns. SIRT1 independent changes are those changes that occur regardless of SIRT1 KO while SIRT1 dependent changes are those that are affected by SIRT1. Note that we consider changes based on the effect size direction allowing clustering based on gene expression pattern.

Figure 9 depicts a clustergram of all SIRT1 independent changes. A top cluster including Creb3l3, Apoc2, Prg4 and Apoa5 shows increased levels with HFD and reduction with NMN in both the Control and SIRT1 KO samples. These genes are members of several enriched pathways including cholesterol metabolism, PPAR signaling and PI3K-Akt signaling and include MCL cluster 1 and 2 members. Other proteins with large changes with a similar pattern include Spta1, Fabp3, Ran and Ldhb. At the opposite extreme are genes that are downregulated with HFD but increased with NMN. These include Serpina1e, CA3, Gstm1, Tnc, and Lifr. These genes are in enriched metabolic and complement and coagulation cascade pathways. Note also the presence of a cluster containing Mup2, Pigr, Hpd, Mup8, Fabp1 and Akr1c6. These genes all have reduced gene expression in HFD but increased expression in response to NMN supplementation. Interestingly they all fall in MCL cluster 2 and have several lipocalin domain proteins (Mup2, Mup8 and Fabp1) and are members of enriched metabolic pathways, tyrosine metabolism, PPAR signaling pathway and fat digestion and absorption.

**Figure 9:**
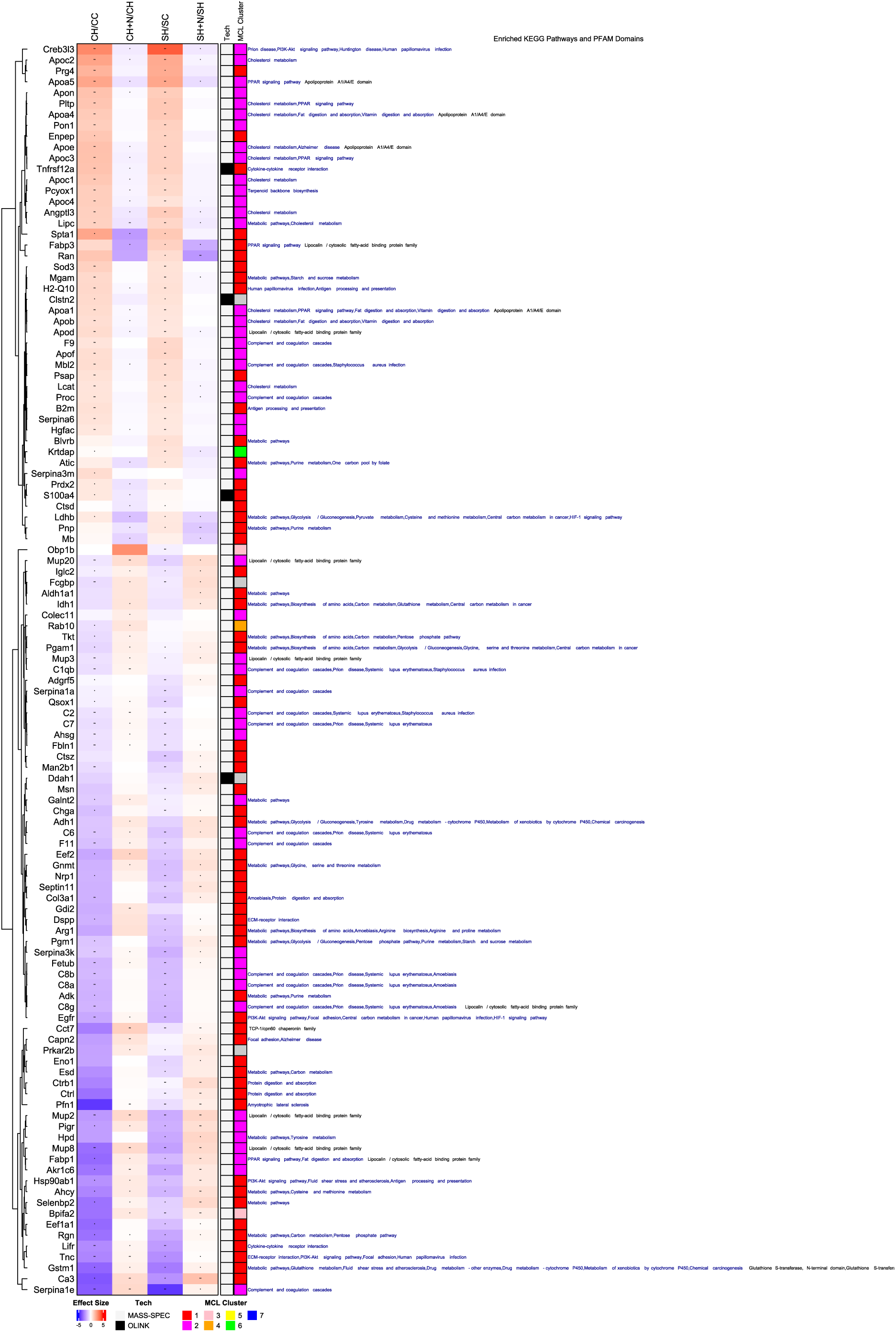
**SIRT1 Independent Diet and NMN effects**. Clustered heatmap of proteins with SIRT1 independent effects in showing the effect size of each comparison, the proteomics technology used (Tech), the MCL cluster and any enriched KEGG pathways (Blue text) and PFAM domains (Black text). The heatmap column names are shortened for clarity and are CC: Control Chow, CH: Control HFD, CH+N, Control HFD + NMN, SC: SIRT1 iKO Chow, SH: SIRT1 iKO HFD, SH+N: SIRT1 iKO + NMN. The asterisks inside each cell indicates significance: * = p-value < 0.05, ** = adj. p-value < 0.05.

In Figure 10, we show SIRT1-dependent changes. Figure 10A shows NMN effects that are impacted by SIRT1 KO. We see that while Saa1 and Saa2 genes (both in MCL Cluster 2) are upregulated in response to HFD, they are slightly reduced with NMN with SIRT1 KO but slightly increased with NMN in the control mice. Two other genes Plin1 and Apeh show upregulation with HFD in the controls and SIRT1 KO but only reduction with NMN in the control. Plin1 is a member of the enriched PPAR signaling pathway while Apeh is not a member of any enriched pathway or any MCL cluster. Figure 10B shows SIRT1 KO genes that impact only the diet effect. Acat3 and Chit1 are significantly increased with HFD in control mice but reduced with HFD in SIRT1 KO mice. Conversely, genes Gfra1, Serpina11, Gm203090, Dnpep, Hspa5 and Ldha are significantly increased with HFD in the SIRT1 KO but are decreased with HFD in the control mice. All these genes are not enriched in any KEGG pathways or PFAM domains. Finally, Figure 10C shows 12 genes that show SIRT1 dependent differences in both diet and NMN. A striking example is that for Il18bp which is increased in HFD in SIRT1 KO and reversed by NMN while the opposite happens in control mice. Ltga6 is another gene with a greatly reduced expression in HFD in control mice which is reversed with NMN but this pattern is slightly reversed in the SIRT1 KO.

**Figure 10:**
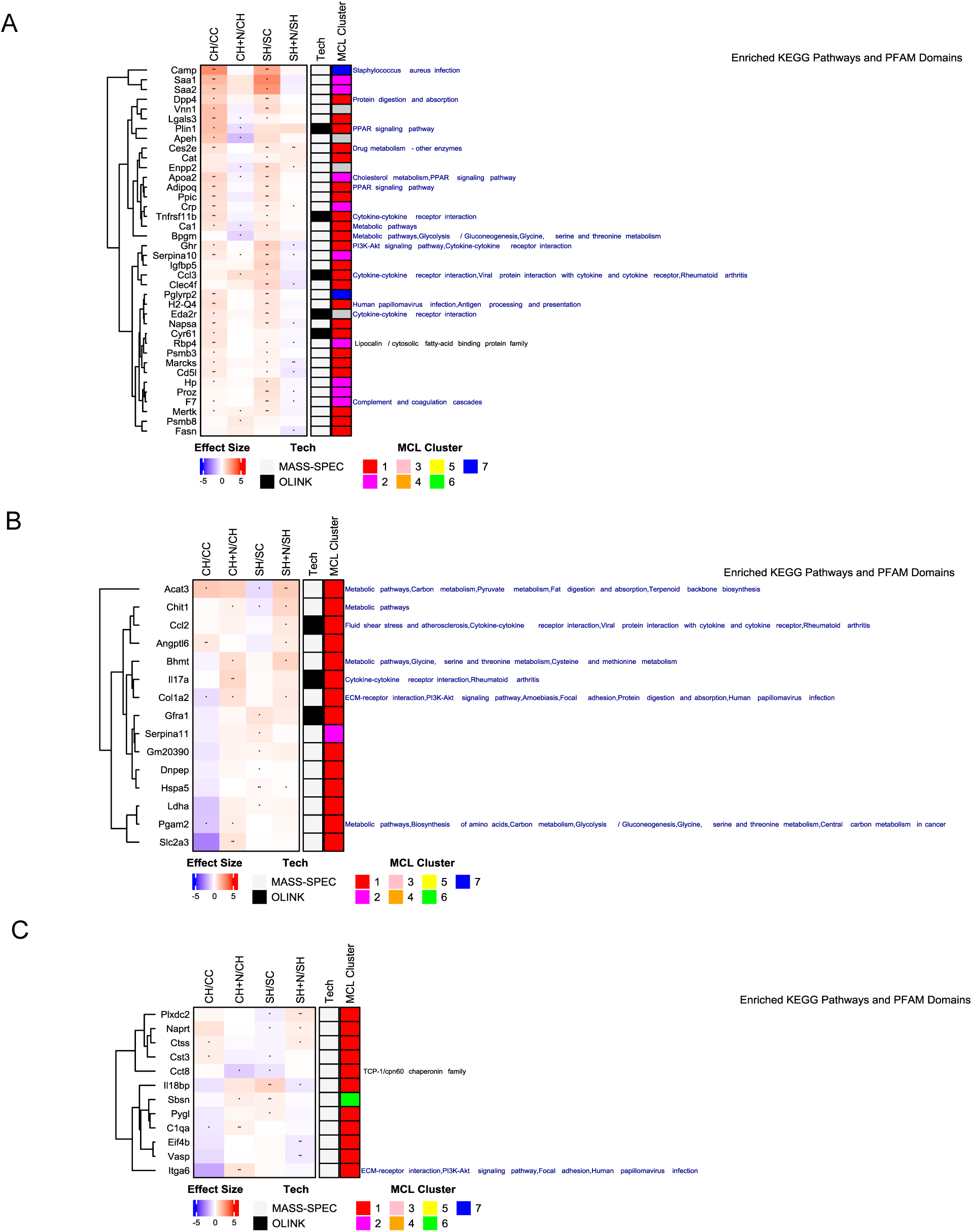
**SIRT1 Dependent Diet and NMN effects**. A) NMN-specific SIRT1 dependent clustered heatmap, B) Diet-specific SIRT1 dependent genes, C) Both diet and NMN SIRT1 dependent genes. Clustered heatmap shows the effect size of each comparison, the proteomics technology used (Tech), the MCL cluster and any enriched KEGG pathways (Blue text) and PFAM domains (Black text). The heatmap column names are shortened for clarity and are CC: Control Chow, CH: Control HFD, CH+N, Control HFD + NMN, SC: SIRT1 iKO Chow, SH: SIRT1 iKO HFD, SH+N: SIRT1 iKO + NMN. The asterisks inside each cell indicates significance: * = p-value < 0.05, ** = adj. p-value < 0.05.

### Causal Analysis of NMN effect on Diet Induced Obesity

To gain greater insight into potential upstream regulators of the observed changes, we performed causal analysis with Ingenuity Pathway Analysis software. Causal analysis attempts to identify regulators where the directionality of the observed expression changes is consistent with the observed effects reported in the literature or in public datasets. We show in Figure 11 the results of this analysis where we aggregate the top 20 master regulators by z-score and by p-value and the expression of their downstream targets present in our data. The analysis reveals a cluster of Fbxw7 and Sirt3 as the most significant activated master regulators, which are linked to molecules including upregulated T_H_17-derived cytokines (Il17f, Il17a) and downregulated metabolism-related proteins (Fabp3, Pdfgb, Ckm and Gsr). In addition, Adam9 and Crmp1 were also identified as upstream regulators that were significant activated. Interestingly, we also observe a cluster of two predicted inhibited master regulators consisting of Adipor2 and Prdm16 linked to a cluster of upregulated Mup proteins. The opioid receptor Oprd1 was also predicted to be inhibited in our analysis (Figure 11). The network between Fbxw7 and its targets is shown in Figure S3 showing dense direct and indirect interactions.

**Figure 11:**
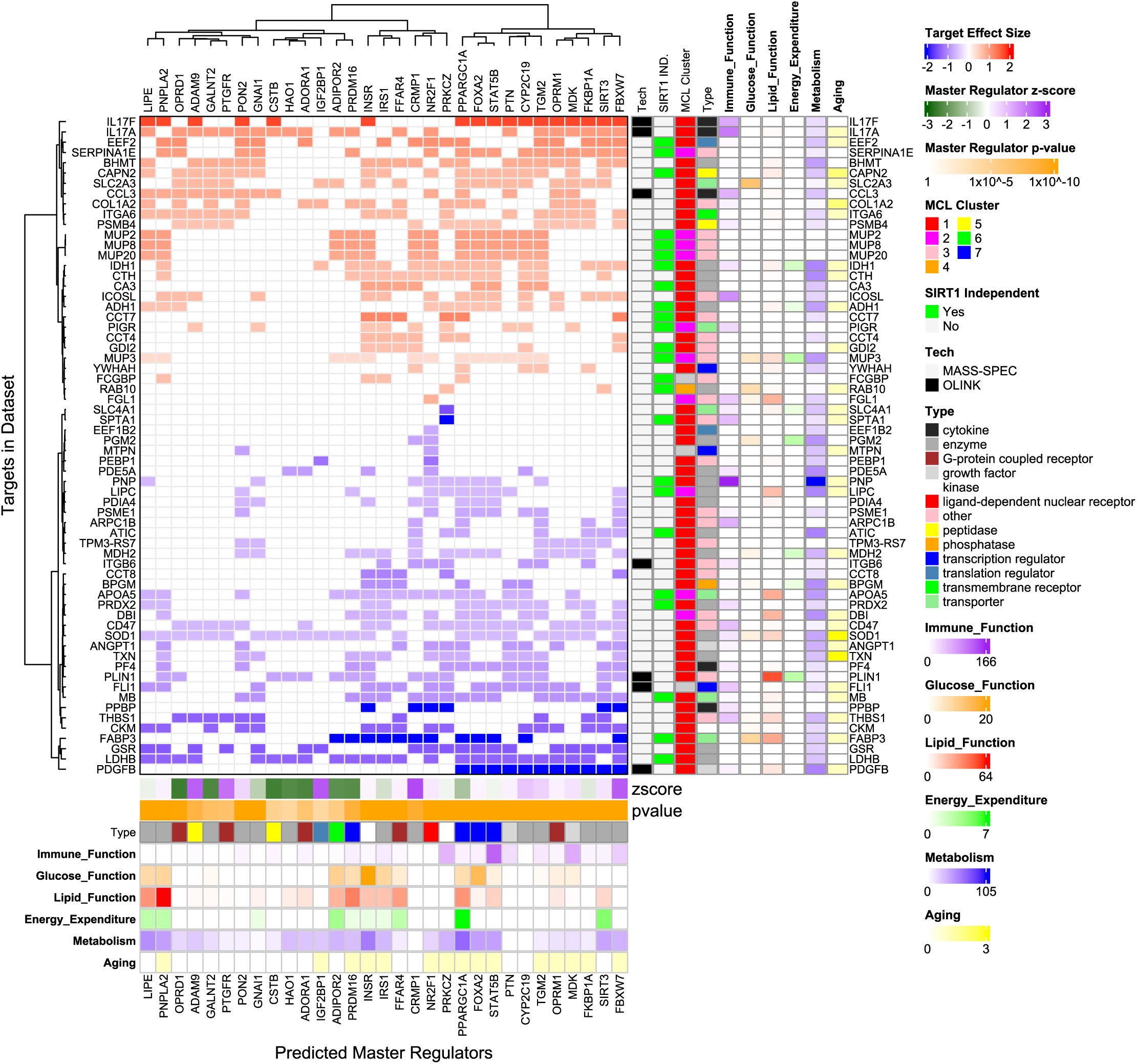
Clustered Heatmap of all Master Regulators with abs(z-score) ≥ 2 and the top 20 most significant by p-value in the NMN + HFD vs. HFD comparison. Different annotations are as indicated in the legend.

In terms of phenotype relevant functional impact, we observe several genes with known association and enrichment in Glucose function (transporter Slc2a3, Mup3, Rab10, enzymes Pgm2 and Sod1 and transporter Fabp3. Slc2a3 and Mup3 are upregulated in response to NMN while the others are downregulated. We also observe genes affect lipid function including peptidase Capn2, enzymes Idh1 and Lipc, Mup3, Fgl1, transporters, Apoa5 and Fabp3 and cytokine Ppbp. A few genes are also associated with energy expenditure including Idh1, Mup3, Slc4a1, Pgm2, Mdh2 and Plin1. Several genes linked with aging or senescence are also observed including Capn2, Col1A2, Sod1 and Txn.

These genes and functions are strongly associated with at least one predicted master regulator. Among the predicted master regulators, we observe a cluster of strongly associated glucose metbaolism genes (Insr, Irs1 and Ffar4). We also observe a lipid associated cluster of enzymes Lipe and Pnpla2. We also observe an energy expenditure linked cluster of transcription regulator genes Ppargc1a, Foxa2 and Stat5B.

The causal analysis also reveals several genes with little to no known association with immune, glucose, lipid, energy or aging functions. Upregulated genes in this category include Eef2, Serpina1e, Mup2, Mup8, Mup20, Fcgbp, Pdia4, Tpm3-rs7 and Mb. From the upstream causal analysis, we observe Crmp1 and Cyp2c29 (closest homolog to Cyp2c19) as genes with little or no known links to our observed phenotypes.

## Discussion

Our data show that oral administration of NMN significantly protected against HFD-associated weight-gain in Control but not SIRT1 iKO mice. Fasting blood glucose levels were elevated in HFD-fed mice, which were significantly normalized in NMN-fed Control but not SIRT1 iKO mice. Importantly, NMN administration enhanced energy expenditure (EE) and RER during periods of activity in Control but not SIRT1 iKO mice. Finally, we report comprehensive plasma proteome changes that were triggered by HFD consumption, NMN administration, and SIRT1 deletion, identifying key genes and pathways involved in key metabolic processes including glucose metabolism, cholesterol metabolism and immune related gene networks. Our discoveries also align favorably with the effects of NMN supplementation in human subjects [55].

### Comparison with previous studies

In order to allow a comparison between our study and the published literature, we selected a dose and route of administration for NMN that is similar to what has been reported previously [38, 56]. Importantly, a protective effect of oral NMN in age-associated weight-gain in chow-fed mice has been reported previously [38]. Because the effect of NMN has been widely investigated in chow-fed mice [57], we focused these studies on obese mice. NMN was well-tolerated by the mice as indicated by similar water intake. Our phenotypic results are broadly consistent with findings of NMN induced weight loss reported in 2023 by Zhang *et al* [39]. The blood proteome results in diet induced obesity are also similar to those reported in Stocks *et al* in ob/ob mice, a model of genetic obesity [15] with increased expression of apolipoproteins and reduction in MUPs and some complement proteins. The resistance to weight-gain in NMN-fed mice is not due to reduced food consumption because food intake was not significantly different between the mice that were fed HFD alone or HFD+NMN. In contrast, SIRT1 iKO mice fed HFD gained less weight than Control mice, and this may be due to a small but significant decrease in food intake. However, the ‘protective’ effect of NMN against weight-gain was not observed in SIRT1 iKO mice. Consistent with previous studies in male mice, we did not observe any significant benefit in glucose-tolerance or insulin-sensitivity, but NMN administration exerted a significant protective effect against dyslipidemia in obese mice, an effect that was again heavily SIRT1-dependent. The SIRT1-dependent increase in EE is consistent with a previous report that showed increased EE in mice that overexpressed SIRT1 [58]. SIRT1 overexpression did not improve skeletal muscle insulin resistance in obese mice [59]. A previous study using nicotinamide also reported data consistent with ours, particularly in terms of a protection against weight-gain, enhanced EE, and a weak or no effect on glucose-tolerance [60].

### Similarity between phenotype and proteome changes

The main significant phenotypes we observed are increased weight, impaired fasting glucose, increased cholesterol and LDL with high fat diet which is reversed with NMN and was associated with increased energy expenditure. SIRT1 iKO reduces the protective effects of NMN consistent with the role of SIRT1 in regulating energy metabolism [61]. These phenotypic changes are mirrored in the proteomic data. First, SIRT1 iKO leads to a significant reduction in plasma proteins relative to control mice for chow, high fat diet and NMN+ high fat diet mice. Second, significant plasma proteins are enriched in key metabolic pathways affecting cholesterol metabolism, PPAR signaling, amino acid and glycolysis pathways. Third, there are SIRT1 independent and dependent genes reflecting the complex SIRT1 iKO phenotype. Fourth, when comparing SIRT1 proteomic effects in the same condition (for example SIRT1 KO CHOW vs Control Chow), we see a large downregulation of most proteins. This is consistent with the SIRT1 KO phenotype of lower weight compared to Control mice (Figure 1A,1C and Figure S1) at the same mouse age in the chow, HFD and HFD+NMN conditions. The weight difference between Control and SIRT1 KO mice on a Chow diet is reflected in relatively few significant proteomic changes compared to Control and SIRT KO mice on a HFD diet and HFD diet+NMN (Figure 4B, Figure 7).

### Key transcription factors in NMN supplementation biology

We find Creb3l3 (aka CREB-H) to be a SIRT1 independent highly increased transcription factor in response to HFD which can be reduced by NMN. Creb3l3 is secreted in the liver and gut in mice and has been shown to be involved in metabolic inflammation [62] and lipid metabolism [63]. We also predict through upstream causal analysis that Pprgc1a (aka PGC-1a) mediates several downstream observed targets including Sod1, Lipc and Fabp3. In the same cluster as Pprgc1 are Foxa2 which has been shown to mediate transcriptional response to fasting [64] and Stat5b linked to immune function [65]. Pprgc1a has been shown to be involved in numerous mitochondrial energy metabolism processes and mice knockouts have shown resistance to high fat diet induced insulin resistance [66].

### Plasma proteomics: cholesterol metabolism and immune related genes

Our proteomic analysis has also highlighted an interesting connection between different genes and pathways. The cholesterol metabolism and coagulation cascade/immunity associated genes that are changed in our analysis especially in the HFD mice are also part of the same MCL Cluster 2 due to co-expression and co-mention in the literature. Our data shows that upregulation of cholesterol metabolism genes is linked to downregulation of many complement/coagulation genes. In addition, causal analysis shows many immune related proteins regulated similarly as genes known to affect lipid function. The complement system mediates many immune functions including defense, inflammation, pathogen targeting and infected cell lysis [67]. Human diseases linked to the complement system include asthma and systemic lupus erythematosus [67]. Deregulation of complement proteins affects immune response [68]. This data suggests that lipid dysregulation is strongly linked with immune function dysregulation which could underly inflammatory diseases like atherosclerosis.

In addition, we observe generally numerous links between proteins as shown in the string-db analysis and in the causal analysis. NMN supplementation of HFD fed mice shows systemic proteomic profile changes.

### Mouse urinary proteins (MUPs) and signaling

We have also observed genes that are in different pathways but contain the same lipocalin binding domain. Lipocalin domains [69] are part of a family of lipid binding transporters [70]. This includes genes that are members of enriched PPARG and cholesterol metabolism pathways and also MUP proteins. In our data, MUP proteins are downregulated in obese mice and upregulated in response to NMN. MUP proteins were originally identified as secreted in mouse urine but have since been found to play a role in nutrient signaling in mice [71, 72]. The highly similar expression pattern for both the known signaling genes (ex. FABP3) and MUP proteins strengthens the idea that MUPs could play a more central role in metabolism and metabolic signaling in mice than previously appreciated.

Interestingly, while humans have not been reported to have functional analogs of mouse MUPs, humans have several lipocalin proteins including ApoD, ApoM and RBP4 which play an intrinsic role in metabolism and LCN2, C8G, ORM1, ORM2 and PAEP which are associated with various immune functions [73]. Many human lipocalin containing proteins have not yet been functionally characterized and it would be interesting to investigate if they play an orthologous role as MUPs in mouse metabolism.

### SIRT1 dependence

When comparing the diet effect and NMN effect in the control and SIRT1 KO based on gene expression direction (Figure 9-10), we observe that while most proteins act in a SIRT1 independent manner, there are several proteins which show clear SIRT1 dependence. Saa1 and Saa2 are two similar proteins which are reversed with NMN in the SIRT1 background but not in the control background. Saa1 and Saa2 are parts of the serum amyloid A family and are expressed in the intestine and liver and play a role in atherosclerosis [74] and cholesterol efflux [75]. Perilipin 1 (Plin1) is another protein with SIRT1 dependence which shows increase with HFD and reversal with NMN in control mice but no significant changes in the SIRT1 background. Plin1 is a lipid-droplet binding protein and plays a role in regulating insulin sensitivity [76].

### Predicted Regulators of NMN effect

Combining our analysis with predictive analysis allows us to build a model of transcription factors that may underlie NMN effects. The most significant predicted activated transcription factor is Fbxw7 as it has a high z-score (indicating observed expression directions are consistent with expected expression direction) and a low p-value (indicating that it affects many proteins). Fbxw7 is part of the ubiquitin ligase system. Specifically, it is a substrate recognition protein that targets for degradation several lipid metabolism genes [77] acting in some proteins through Notch2 [78]. Two other interesting inhibited potential regulators are Adiponectin receptor 2 (Adipor2) and PR domain containing 16 (Prdm16). In our analysis, Adipor2 is predicted to be inhibited in response to NMN in HFD mice. A previous study showed that knocking out Adipor2 in mice led to improved dyslipidemia but worse glucose control [79], consistent with some of the observed effects in NMN supplementation on HFD mice. Prdm16 is an extensively studied gene with observed effects in thermogenesis [80] and exercise induced adipose tissue remodeling [81]. Both Adipor2 and Prdm16 are linked to the expression of MUP proteins. Our analysis also identified predicted inhibition of the opioid receptor Δ (Oprd1), which is interesting for two reasons. First, because Oprd1 KO mice show enhanced energy expenditure (despite being hyperphagic), thereby resisting weight-gain when fed HFD [82]. Second, because variants in Oprd1 have been previously linked to adiposity and glycemic control [83]. Our analysis also predicted activation of Crmp1, which is interesting considering its link to age-related diabetes [84].

### Limitations

Our study provides strong evidence for the beneficial effects of NMN supplementation in obese mice, including improving dyslipidemia, reducing weight-gain, and acting through well-established metabolic and immune related pathways. However, further investigations are required to address some of the limitations of our study. One limitation is that our proteomic data comes from blood, so the tissue origin of the secreted molecule is not clear. Second, direct investigations of protein activity would be valuable in assessing if the altered expression levels correlate with altered function. Third, while our data support the predicted causal effects, direct evidence of causality requires additional experiments. Mechanistic inference is further complicated by the fact that we measure the proteins in blood while the effects and protein levels could be different in different tissues.

Overall, our results show a protective effect of NMN on diet induced obesity in mice which is partially mediated by SIRT1. These phenotypes are reflected in blood proteome changes in linked lipid metabolism and immune pathways and through different metabolic pathways.

## Data availability

All the datasets generated in the current study are available from the corresponding author upon request.

## Author contribution

YM designed and performed experiments, analysed and interpreted data, and co-wrote the manuscript. NMH analyzed the proteomics datasets, performed statistical analysis, and co-wrote the manuscript. RE, HS, MNA, and SC performed the proteomics experiments, analyzed data, and contributed to manuscript preparation. MVA, AYM, and MV performed experiments. FS supervised the proteomics experiments, analysed and interpreted data, and contributed to manuscript preparation. NAM supervised the project and assisted with data interpretation and manuscript preparation. All the authors read the manuscript and approved its submission.

## Conflict of interest

The other authors declare that they have no conflicts of interest.

Ethics approval: The Institutional Animal Care and Use Committee (IACUC) at Weill Cornell Medicine-Qatar approved all the animal experiments (Protocol 2015-0026) and the project was carried out in an AAALAC International accredited facility.

## Acknowledgements

The authors acknowledge support from the proteomics and bioinformatics core facilities at Weill Cornell Medicine-Qatar. We are grateful to Ms. Cindy Conti and her team from the WCM-Q vivarium core for their assistance with the animal experiments. The core facilities are supported by a Biomedical Research Program (BMRP) grant awarded by Qatar Foundation. We are grateful to Dr. David Sinclair for providing the SIRT1 transgenic mice. We also acknowledge Drs. Michael Bonkowski (RIP) and Alice Kane from the Sinclair lab for assistance with shipping NMN to Qatar, and Juliette Wipf and her team from the Laboratory of Comparative Pathology (LCP) at Memorial Sloan Kettering Cancer Center (MSKCC) for performing the plasma lipid analysis. The LCP is supported by a P30 core grant.

---

**Figure S1:**
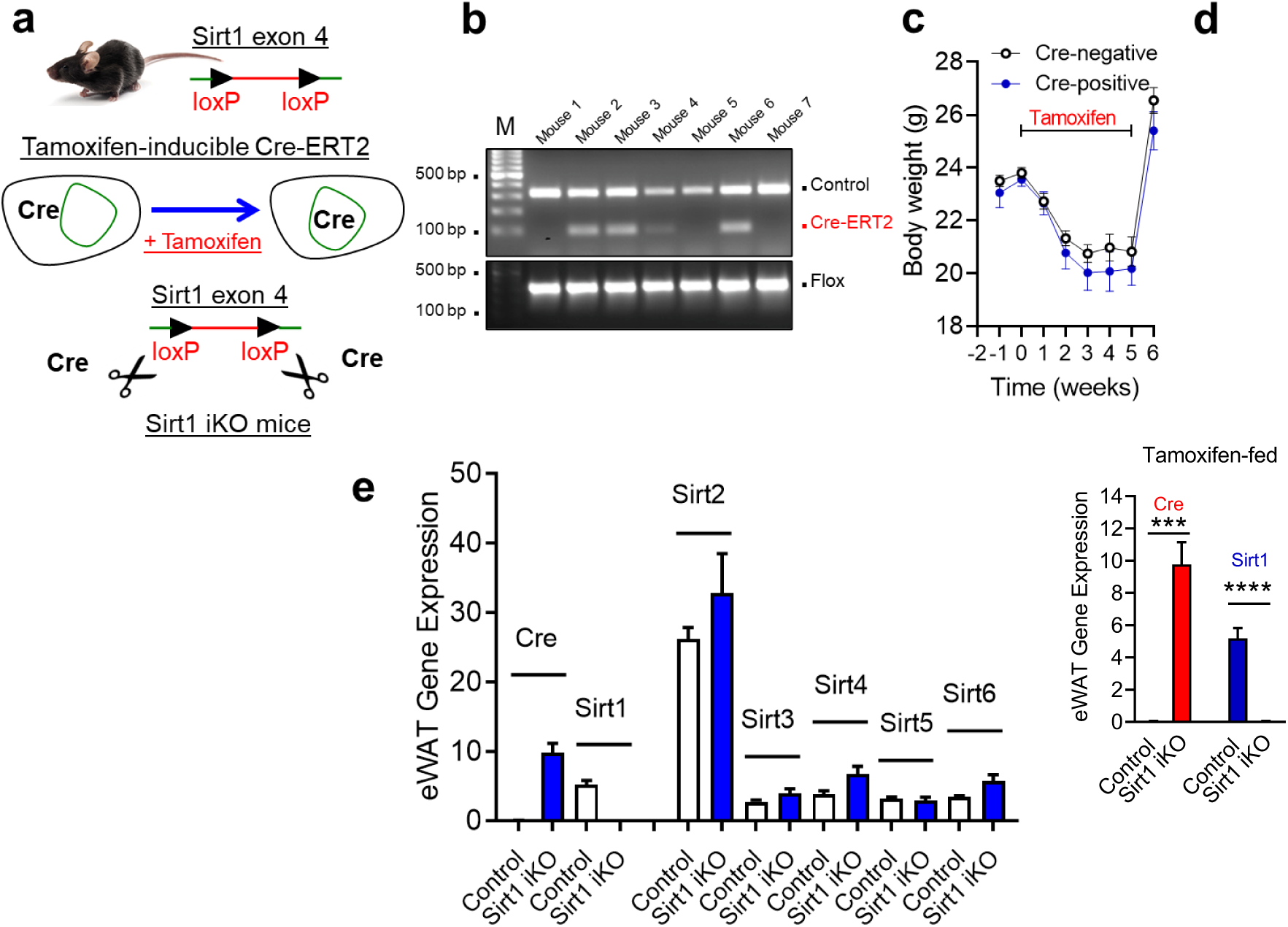
Inducible and specific deletion of SIRT1 in mice. (a) SIRT1-deletion strategy using the tamoxifen-inducible Cre-ERT2 system. Cre-recombination leads to the deletion of exon 4, which encodes the catalytic site in SIRT1. (b) Genotyping to identify Cre-negative and Cre-positive mice. Male Cre-positive mice were bred with Cre-negative female mice to generate Cre-positive and – negative mice in the same litter. The data are shown for mice prior to tamoxifen administration. (c) Weight loss in tamoxifen-fed mice and recovery after tamoxifen withdrawal. (d) qPCR data showing the expression levels of Cre and SIRT1 in the indicated groups. (e) qPCR data showing a specific loss of SIRT1 mRNA in eWAT after tamoxifen administration.

**Figure S2:**
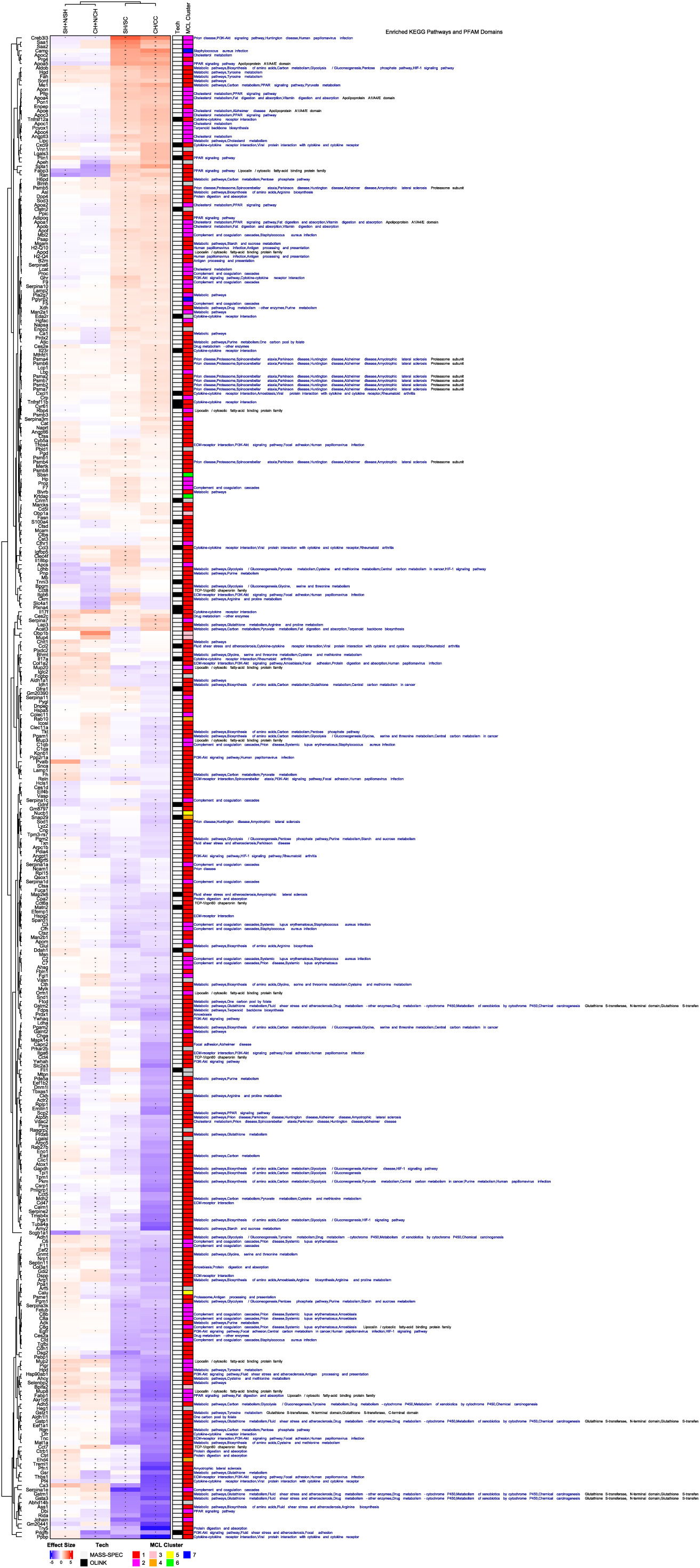
Grouped Analysis Gene Expression Heatmap. Clustergram shows the row and column clustered protein expression of all proteins in four differential expression comparisons. The row annotations are the Tech (Mass-spec or Olink) and the MCL cluster the protein belongs to.

**Figure S3:**
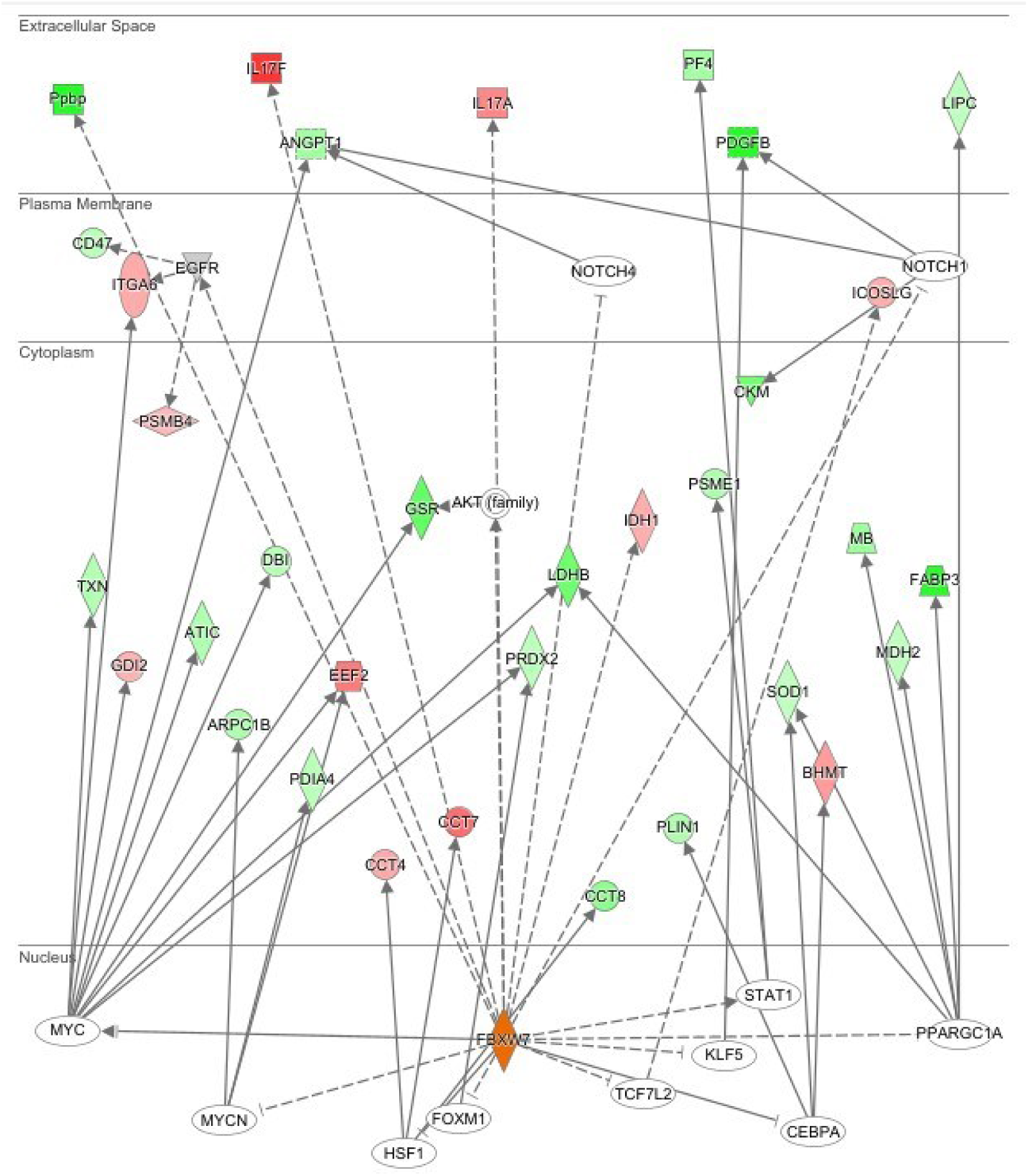
**FBXW7 Network in the NMN + HFD vs. HFD comparison**. Node colors represent expression with red increased and green decreased. Solid and dashed lines are direct and indirect interactions. FBXW7 is highlighted in orange.

